# Retrograde Control of Cytosolic Translation Targets Synthesis of Plastid Proteins and Nuclear Responses for High-Light Acclimation

**DOI:** 10.1101/2021.02.18.431817

**Authors:** Marten Moore, Aaron B. Smith, Melanie Wegener, Corinna Wesemann, Sonja Schmidtpott, Muhammad Ansar Farooq, Diep Ray Ganguly, Thorsten Seidel, Barry J. Pogson, Karl-Josef Dietz

## Abstract

Canonical retrograde signalling comprises information transmission from organelles to the nucleus and in particular controls gene expression for organellar proteins. The need to re-assess this paradigm was suggested by discrepancies between de novo protein synthesis and transcript abundance in response to excess light. Here we uncover major components of a translation-dependent retrograde signalling pathway that first impacts translation and then transcription. The response realization depends on the kinases Mitogen-activated protein kinase 6 (MPK6) and Sucrose non-fermenting 1-related kinase (SnRK1) subunit, AKIN10. Global ribosome foot-printing revealed differential ribosome association of 951 transcripts within 10 min after transfer from low to high light. Despite predominant translational repression, 15 % of transcripts were increased in translation and enriched for chloroplast-localized photosynthetic proteins. About one third of these transcripts, including Stress associated proteins (SAP) 2 and 3, share regulatory motifs in their 5′-UTR that act as binding sites for glyceraldehyde-3-phosphate dehydrogenase (GAPC) and light responsive RNA binding proteins (RBPs). SAP2 and 3 are both translationally regulated and interact with the calcium sensor Calmodulin-like 49 (CML49), which promotes relocation to the nucleus inducing a translation-dependent nuclear stress response. Thus, translation-dependent retrograde signalling bifurcates to directly regulate a translational circuit of chloroplast proteins and simultaneously initiate a nuclear circuit synchronizing retrograde and anterograde response pathways, serving as a rapid mechanism for functional acclimation of the chloroplast.

## Introduction

Light impacts plant morphogenesis, development, and metabolism. Gains in biomass and fitness strongly correlate with light-dependent photosynthesis and starch accumulating during the light phase being essential for metabolism and growth during the dark phase. However, excess or high light (HL) leads to an over-reduction of the photosynthetic electron transport (PET) chain and enhances production of reactive oxygen species (ROS) [1]. The natural light environment fluctuates continually on time scales of seconds, hours, days, and seasons. To cope with rapid light fluctuations, plants control their capacity to dissipate excess energy and minimize photo-oxidative damage [2]. ROS produced in excess light are quickly detoxified, for example H_2_O_2_ by the water-water cycle in an ascorbate-dependent or ascorbate-independent process utilizing ascorbate peroxidases and peroxiredoxins, respectively [3, 4]. On longer time scales, changes in light quantity alter photosystem stoichiometry and increase the photosystem repair capacity [5].

Proteins of the photosynthetic machinery are encoded in the plastid and the nuclear genomes while all antioxidant enzymes are exclusively encoded in the nucleus, as is the majority of plastidial proteins. Transcription, stability, and translation of chloroplast-encoded transcripts strongly depends on nuclear-encoded sigma factors and RNA binding proteins (RBPs) [6]. The reorganization of photosystems and readjustment of the antioxidant systems, therefore, rely on a coordinated synthesis of nucleus- and chloroplast-encoded proteins [7]. Coordination is realized by transmitting information on the metabolic state of the chloroplast as retrograde signals to the nucleus. Alterations in light quality and quantity result in fluctuations of metabolites and signal export from the chloroplast with impacts on transcript abundance within minutes [8–10].

As the majority of chloroplast proteins are nuclear-encoded, several signaling pathways exist for developmental and operational retrograde control of nuclear gene expression. They include signals from the plastoquinone and carotenoid pools, tetrapyrrole biosynthesis, dihydroxyacetone phosphate (DHAP), ROS, ascorbate/dehydroascorbate (Asc/DHA) and glutathione/glutathione disulphide (GSH/GSSG), hormones like 12-oxo-phytodienoic acid (OPDA) that also functions as precursor for jasmonic acid (JA), and xanthoxin as precursor of abscisic acid (ABA), the SAL1/PAP pathway and sugars [3, 9, 11–18]. Rapid light induced transcription requires export of triose phosphates, such as DHAP, via the Triose phosphate/phosphate translocator (TPT2) and Mitogen activated protein kinase 6 (MPK6), which is coupled to the upregulation of a subset of Apetala 2/Ethylene response factor (AP2/ERF) transcription factors [9]. It is worth noting that TPT2 co-controls the transmission of the DHAP retrograde signal [19].

To date, retrograde signaling pathways are considered with respect to nuclear gene regulation. However, direct retrograde control of cytosolic translation remains mostly unexplored, especially during rapid responses to stimuli. A few studies focused on the effects of translational control under the effects of the controversially discussed gun mutants [20], or on targeted transcripts without whole genome analysis [21, 22], which allow a limited scope of regulative mechanism of cytosolic translation as a consequence of retrograde signalling.

In addition to regulation by phosphorylation/dephosphorylation, redox regulation of the translational apparatus has been suggested as one such translational control mechanism [23]. Recently, light-induced ROS production has been linked to rapid activation of eIF2α kinase General control non derepressable-2 kinase (GCN2), thus inhibiting translation initiation [24]. This process also affected the translation of transcripts encoding mitochondrial ATP synthesis and chloroplast thylakoid proteins.

Post-transcriptional regulation of translation and mRNA stability is known from studies on circadian rhythmicity and involves light entrainment [25]. Transcripts encoding enzymes of central metabolism associate with the ribosomes in in a light- and circadian-dependent manner. The impact of circadian rhythmicity on the steady-state pool of polysome-recruited transcripts was lost if the shift to darkness or re-illumination occurred out of phase [26]. The energy sensor kinase AKIN10 entrains the circadian clock depending on light input [27]. AKIN10 is part of the Sucrose non-fermenting 1-related kinase 1 (SnRK1) complex and acts as a central regulator of transcription and translation networks via the Target of rapamycin (TOR) complex [28–30].

In addition to temporal retrograde signaling cascades affecting translational regulation, consideration to spatial effects should be given. Photosynthetic cells in flowering plants usually have multiple chloroplasts but a single nucleus. The distance between individual chloroplasts and the nucleus often spans several tens of micrometers, potentiating heterogeneity of nuclear gene regulation between chloroplasts of varying vicinity to the nucleus. However, each chloroplast has a surrounding cytosol with translation-competent ribosomes. Retrograde control of translation may be a potent mechanism for immediate spatial and temporal adjustment of the chloroplast proteome. Therefore, there is a fundamental need to address the significance and mechanism of translational regulation of light acclimation by retrograde signalling.

Here we applied ribosome foot-printing to measure global protein synthesis in arabidopsis leaves following a 10 min exposure to HL. Results revealed a rapid light-induced reorganization of the translatome, which was perturbed in *mpk6* and *akin10* mutants. A cluster of transcripts were preferentially polysome-associated in HL and encoded primarily for cytosolic proteins involved in translational and chloroplast-targeted proteins involved in assembly, function, and maintenance of the chloroplast. A second cluster revealed simultaneous increases in transcript abundance and polysome association, containing stress-associated transcripts such as *HSF2A*, *DREB2A*, and *HSPs*. Transcripts of these clusters contained conserved sequence motifs in their 5’-UTRs, which was found to increase translation *in vivo*. Focusing on one motif, we identified light-dependent recognition by GAPC and scrutinized the downstream effects of the translational reorganization following two selected transcripts *SAP2* and *SAP3*, whose proteins interact with the calcium sensor CML49 co-regulating further stress responses. The study evidences the effects and identifies regulators of rapid retrograde regulation of cytosolic translation.

## Results

### Rapid retrograde signaling for translational reprogramming

To investigate the impacts of severe light stress on translation, we combined ribosome profiling with an established light shift protocol to compare differing fold-increases in light intensity on cumulative translation (Figure 1) (*9, 31, 32*). Plants were transferred from normal growth light (NL, 80 µmol photons m^−2^ s^−1^) or low light (LL, 8 µmol photons m^−2^ s^−1^ for 10 d) to HL (800 µmol photons m^−2^ s^−1^, NL➔HL or LL➔HL) representing a 10-fold or 100-fold radiation increase, respectively. Transcripts associated with multiple ribosomes (polysomes) were isolated as described [31] under presence of translational inhibitors cyclohexamide, chloroamphenicol and lincomycin for combinatorial arrest of cytosolic and plastidial ribosomes. Extracts were loaded and run through sucrose gradients, and polysomal profiles recorded during fractionation (Figure 1a). This revealed rapid reorganization of ribosome-associated RNA within 10 min of a 100-fold light increase. Comparison of LL control with 10 and 60 min LL➔HL treatments revealed cumulative dynamic changes in ribosome-associated polysome profiles (Figure 1a). The 80S monosome peak (one ribosome per mRNA) was greatly reduced after 10 min of LL➔HL, but less so by 60 min LL➔HL. The polysomal area was unchanged between LL control and 10 min LL➔HL, but increased after 60 min LL➔HL. Contrarily to the 100-fold light shift, no time- and dose-dependent changes in the polysomal profiles could be observed in 10-fold NL➔HL and NL➔HL➔NL shifted plants.

**Figure 1:**
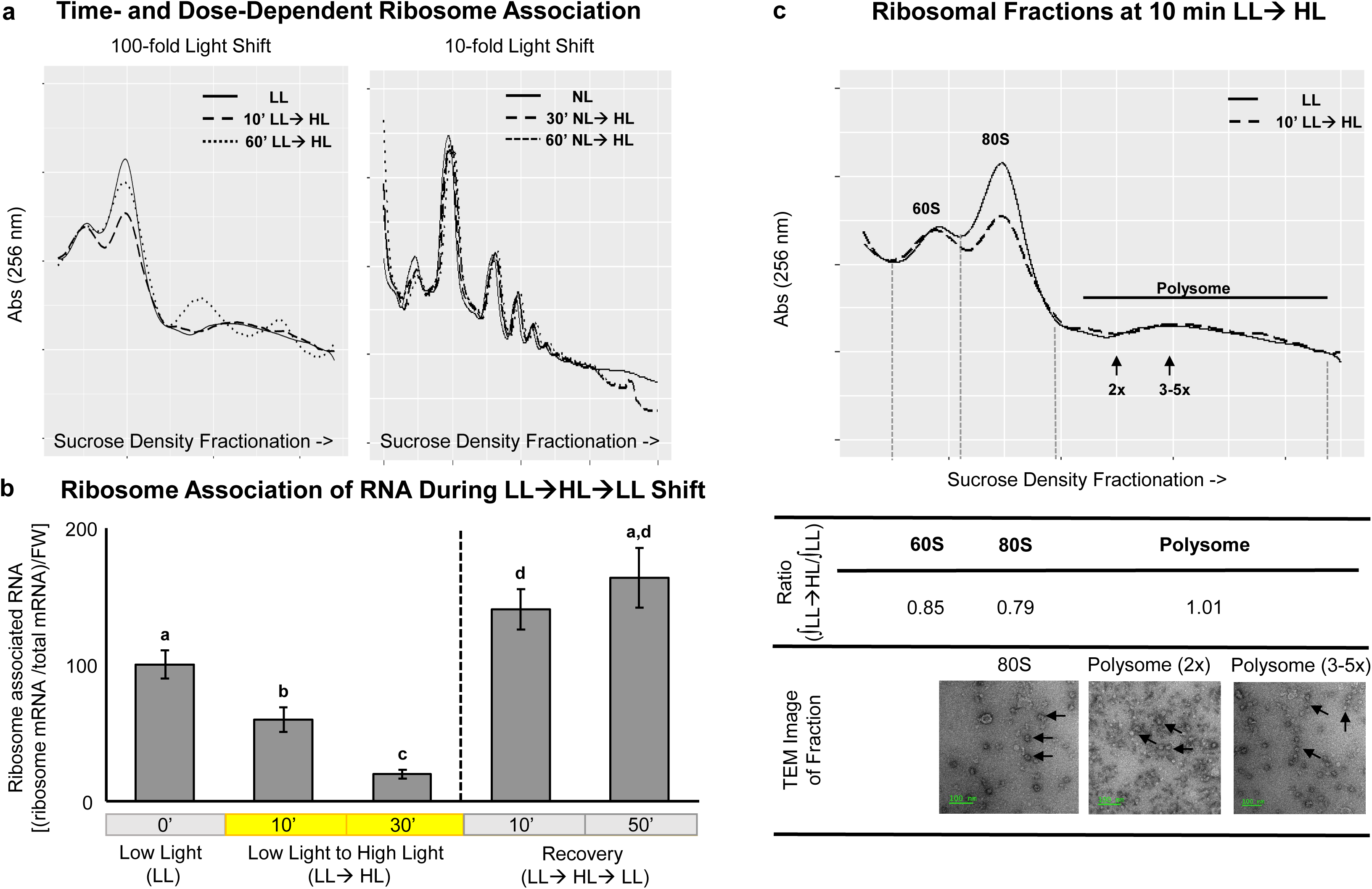
Time- and dose-dependent changes in ribosome associated transcripts during high light treatment. (a) Growth light (NL) or low light (LL)-acclimated plants were transferred to high light (HL) for a 10-fold light shift (NL➔HL) or a 100-fold light shift (LL➔HL) for indicated time. Depicted are representative baseline-normalised polysomal profiles in sucrose gradients of n=3 biological replicates. (b) Association of RNA with polysomes is dynamically and reversibly regulated during a 100-fold light shift. Plants were kept in LL (0‘) or transferred to HL (LL➔HL) for indicated time periods and then transferred back to LL for the indicated time for recovery. Ribosome association was determined by comparing ribosome-associated mRNA quantity from polysomal fractions of sucrose gradients to total mRNA on a fresh weight (FW) basis (mean ± SE from n=3 independent experiments; letters indicate significance of difference as determined with ANOVA post-hoc Tukey p<0.5). (c) Analysis of polysome status from LL and 10 min LL➔HL treated plants via polysomal profiling and TEM imaging of ribosomes (arrows) of the corresponding fractions revealed presence of monosomes (80S), low molecular polysomes (2x) and high molecular polysomes (3-5x) (green bar represents 100 nm).

Quantification of ribosome-associated mRNA isolated from ribosomal fractions in Figure 1a revealed a progressive repression of polysome association during LL➔HL up to 30 min (Figure 1b). To confirm reversibility of the repression, we also quantified ribosome-associated mRNA during recovery from 10 and 50 minutes light stress (i.e. LL➔HL➔LL). We observe equally rapid re-initiation of translation represented by increased ribosome-associated mRNAs within 10 minutes of transferal back to LL following either 10 or 50 min LL➔HL (LL➔HL➔LL). This indicated that the dynamic regulation is rapidly initiated within 10 min as first timepoint of significance, which is expressed in a 0.79-fold decrease in 80S (monosomes) peak area and 0.85-fold decrease in 60S peak area (Figure 1c), resulting in a cumulative down regulation of ribosome-associated mRNA. Ribosome integrity was explored by staining fractions of monosomes, low molecular polysomal mass (2 ribosomes) and high polysomal mass (≥3 ribosomes) and imaging via transmission electron microscopy (TEM). Low and high molecular mass polysomes could be imaged, confirming successful separation of intact monosomes and polysomes.

### Regulatory pathways connect retrograde signals and cytosolic translation

To investigate the upstream regulation leading to the dynamic shift in cytosolic translation, we queried two potential interactors of translational regulation that operate in the same rapid time frame. Previously, we demonstrated phosphorylation of MPK6 in response to retrograde signaling within less than 10 min following the same LL➔HL shift treatment [9]. Intriguingly, MPK6 activity is potentially linked to phosphorylation state of the SnRK1 subunit AKIN10 [29, 32], which acts as negative feedback loop to TOR signaling and the initiation of translation [33, 34]. Therefore, we hypothesized a link between retrograde signaling and translation, through light-responsive activation of MPK6 and AKIN10 [29]. To test this, we compared changes of ribosome profiles in *mpk6* and *akin10* and Col-0 wild-type (WT) background after 10 min LL➔HL shift (Figure 2). Under LL control conditions *mpk6* revealed a mildly reduced 60S and 80S peak area in comparison to WT control (0.91-fold for each) and a 33% reduction in polysome-bound mRNA (Figure 2a). Contrarily, *akin10* exhibited a strong reduction of the 60S and 80S peak area in comparison to WT (0.67 and 0.52-fold, respectively) with no significant difference in polysomal area. After 10 min LL➔HL, *mpk6* exhibited a closer to WT ribosomal profile, but with decreases in 60S and 80S area of 45% and 49%, respectively (compared to 15% and 21% in WT) (Figure 2b). In addition, a strong decline of 43% in polysomal area was recorded. The profiles of *akin10* did not differ between LL and LL➔HL, but regardless of experimental conditions exhibited a lower 60S and 80S peak area in comparison to WT with an unaltered level of polysomal area.

**Figure 2:**
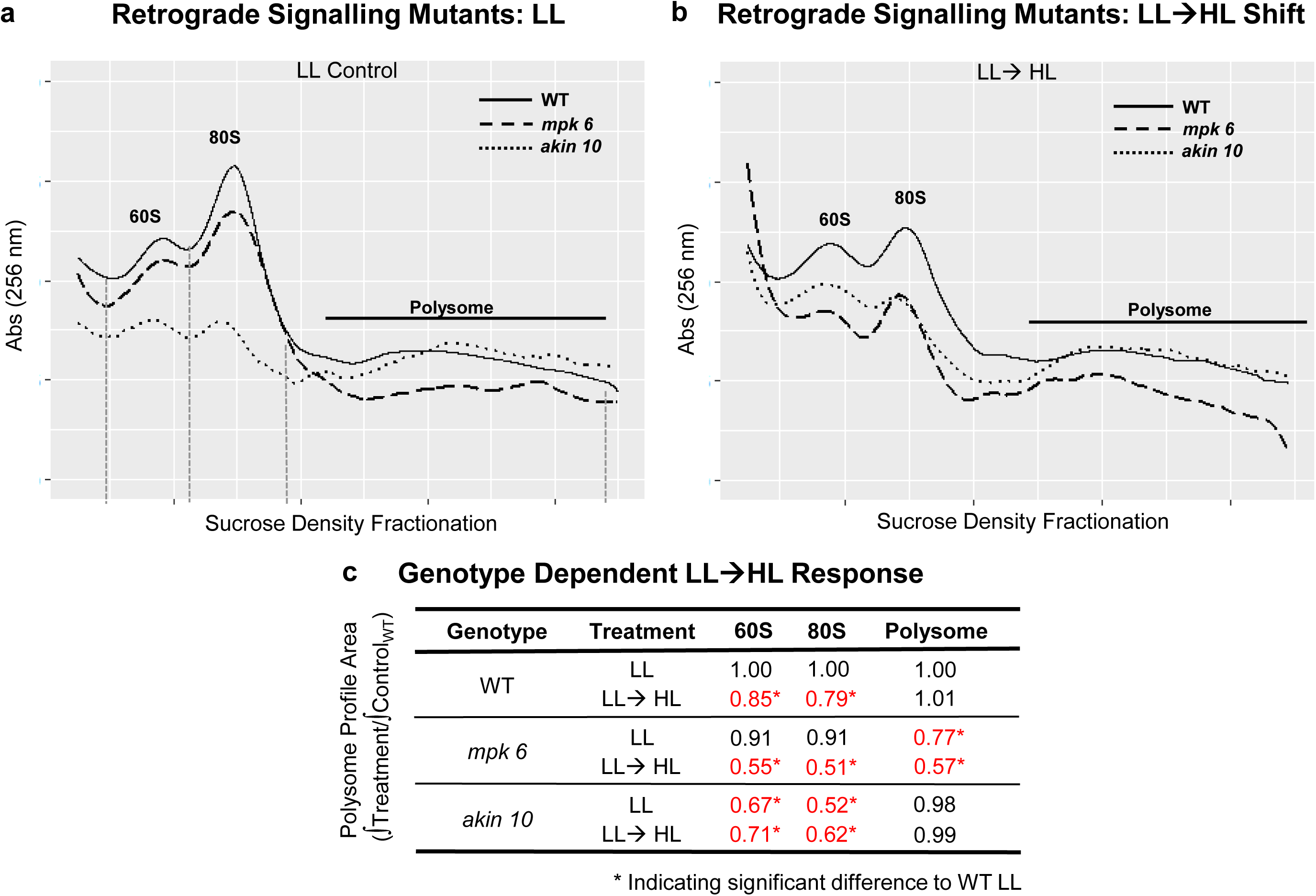
Mutants involved in retrograde signalling exhibit genotype-dependent LL➔HL translational response. (a). Differential ribosome association of transcripts in retrograde mutants as detected in polysomal profiles of LL grown and 10 min LL➔HL treated plants (b). 10 min LL➔HL shift repressed ribosome association of transcripts. Depicted are representative polysomal profiles of WT, *mpk6* and *akin10* KO background after sucrose gradient centrifugation of n=3 biological replicates (60S, 80S and polysomal fractions indicated). (c) Quantification of genotype dependent LL➔HL response shows repression of ribosome association in WT, a stronger decrease in *mpk6* and a loss of regulation on akin*10* background. Polysome profiles were background normalized and the peaks corresponding to 60S, 80S and polysomal area integrated and normalized to WT control (* represents significant differences to WT, two sided Student’s t-test p<0.05).

In conclusion, the loss of MPK6 and SnRK1 subunit AKIN10 showed alterations to different aspects of polysomal profiles that were genotype- and light-dependent, cumulatively altering the translational profile. While *akin10* lost the WT reaction to LL➔HL treatment, *mpk6* reflected the WT reaction in regards of 60S and 80S area, with an additional reduction of polysomal area after LL➔HL treatment.

### Regulation of translation outweighs transcriptional changes in early light acclimation

We quantified the cumulative changes in translation in response to LL➔HL regulation by RNA sequencing on polysome-associated transcripts. Ribosome footprints (RPF) were generated from three independent experiments of LL and LL➔HL-treated plants. RNase If treatment digested the unprotected sequence stretches [35]. Both the RPF (translatome) and total purified RNA (transcriptome) were quantified by ribosomal RNA-depleted RNA sequencing [36]. The generation of footprints in the presence of CHX, CHL and LIN resulted in oligonucleotide read lengths with two local optima of 24-26 (footprint size from 70S ribosomes in plastids and mitochondria) and 28-34 nt lengths (cytosolic footprint size from 80S ribosomes), documenting sufficient capture of arrested plastidial and cytosolic footprints (Suppl. Fig 1a). The read frame periodicity of the footprints (Suppl. Fig 1b), as well as the read frame at start and end sites that corresponded to independent studies using a combination of CHX and CHL for translation inhibition [37, 38].

Performing differential gene expression analysis on ribosome-footprints confirmed the observation of global repression of translation with 793 transcripts (76 % of total DEGs) that were significantly down regulated. A set of 92 transcripts (9 %) remained unchanged and abundance of 158 (15 %) increased (Suppl. Fig 2a). Conversely, total RNA remained mostly unchanged with only 121 and 83 transcripts being transcriptionally up- and down-regulated, respectively. Based on our analysis 4.8-fold more transcripts were regulated at the translational than transcriptional level after 10 min LL➔HL.

### Evidence for a translational retrograde circuit enriched in photosynthesis proteins

In order to reveal distinct regulative patterns not following the cumulative translational response, we further investigated the 24 % of transcripts showing significant ribosome association after LL➔HL. The transcripts showing significantly differential expression in the ribosome-bound or free mRNA were examined for cluster formation by plotting their relative change in ribosome-association versus the change in total amount (FC ribosomal mRNA LL➔HL/LL / FC total mRNA LL➔HL/LL). This evaluation allowed for tentative distinction of four subsets of transcripts with variant regulatory responses (Figure 3a). Transcripts in Cluster 1 revealed increased ribosome association without changes in total mRNA abundance (119 DEGs). Clusters 2 and 3 comprise transcripts that upon transfer to HL simultaneously changed in the ribosome-associated and total mRNA, 62 up and 123 down regulated DEGs, respectively (Figure 3a; clusters 2 and 3). Association with ribosomes of Cluster 4 transcripts decreased in HL, but were not significantly altered in total RNA fold change (FC) (466 DEGs).

**Figure 3:**
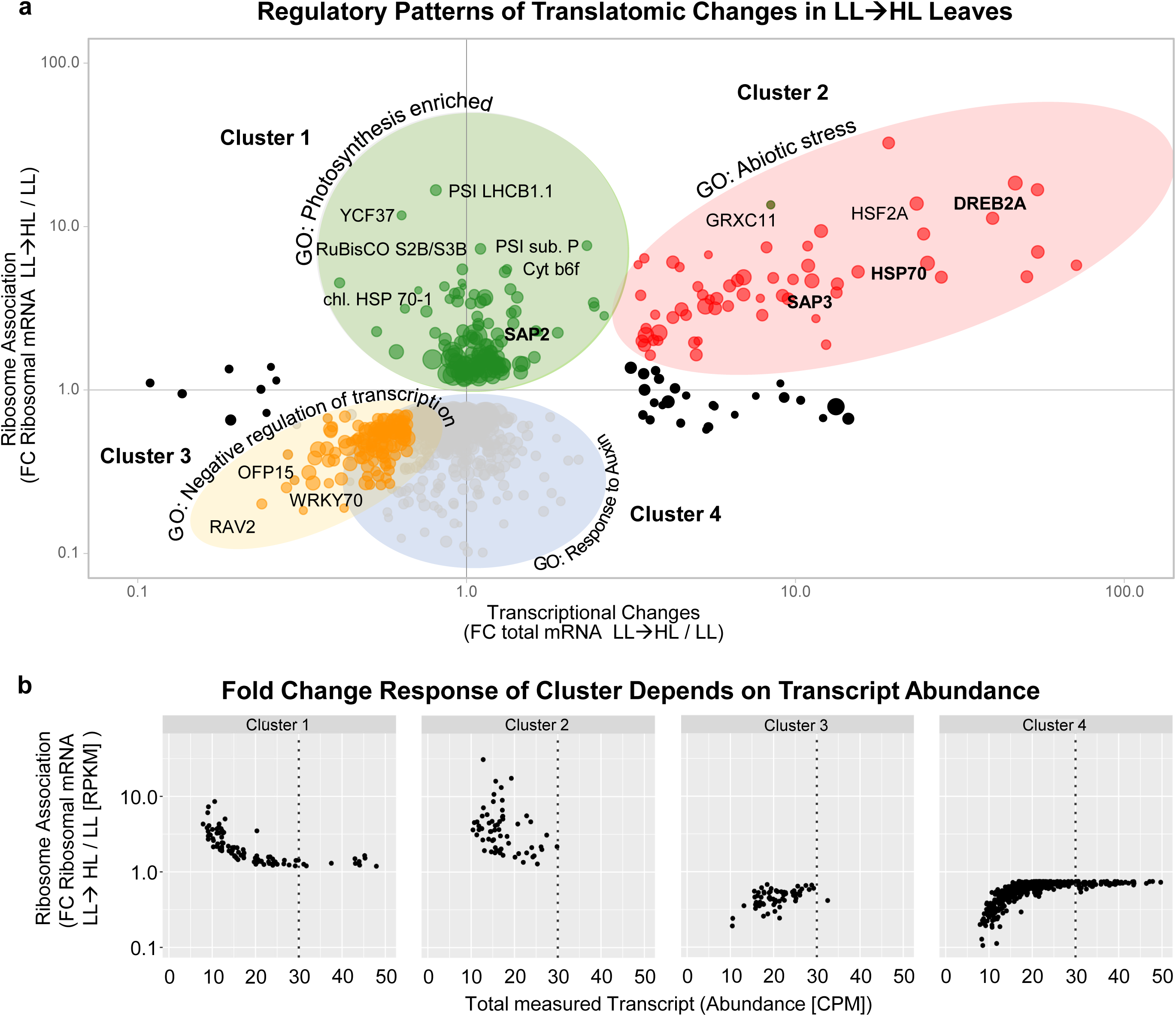
Translatome after 10 min LL➔HL reveals distinct regulatory pattern. Significantly regulated transcripts are depicted as dots with the size indicating transcript abundance. There are 4 distinct clusters of transcripts: translationally upregulated (Cluster 1), transcriptionally and translationally upregulated (Cluster 2), transcriptionally and translationally downregulated (Cluster 3), and translationally downregulated (Cluster 4). Each cluster is significantly enriched in transcripts coding for different GO annotations, exemplar genes are listed. Interestingly, Cluster 2 codes only for stress genes (most prominently HSF2A, DREB2A) while Cluster 1 encompasses almost exclusively photosynthetic genes and translation processes (e.g. LHCBG1.1 and R40S_S16-1). (b) The fold change response of each cluster depends on transcript abundance. Comparison of transcript ribosomal mRNA fold change with total measured transcripts revealed an inverse correlation for Cluster 1 and 4, although they are not transcriptionally changed. Surprisingly, this inverse correlation was not clearly seen for the transcriptionally and translationally changed transcripts in Cluster 2 and 3, indicating a less pronounced impact of transcript abundance (i.e. availability) on ribosome association.

Cluster 1 transcripts that were solely increased in ribosome association were five-fold enriched in transcripts coding for chloroplast localized proteins according to SUBA4 [39]. GO analysis of cluster 1 revealed a 22.9-fold enrichment for the GO term ‘photosynthesis, light reaction’ in comparison to the whole genome (FDR p<4.32E-07). It encompassed transcripts coding in particular for photosystem light harvesting proteins, including minor (AT3G47470) and major light harvesting proteins (AT4G10340), subunits of photosystem I (AT2G46820), one of the two nuclear encoded cytochrome b6f subunits (AT2G26500) and NADPH oxidoreductase subunits participating in electron transport (AT1G06690) as well as small subunits of ribulose bisphosphate carboxylase oxygenase (RuBisCO, AT5G38410) (see Suppl. Dataset 1, cluster 1). In addition to photosynthetic proteins, this set included transcripts for proteins participating in assembly, damage, and repair. For example chloroplast localized HSC70-1 (AT5G02500), which is involved in protein import; chloroplast vesiculation factors (AT4G32150) participating in partitioning damaged proteins for proteolysis; and PYG7/YCF37 (AT2G23670) which is required for PSI assembly/repair. The transcripts coding for cytosolic proteins were 9.36-fold enriched in GO terms for translation (FDR p<4.19E-07), e.g., proteins of ribosomal large subunit (AT3G08520).

Transcripts of cluster 2 with increased ribosome-associated and total mRNA (Figure 3a), were enriched in ‘abiotic stress’-related GO terms, namely response to heat (71.25-fold enriched in comparison to genome, FDR p< 4.53E-41); response to hydrogen peroxide (94.93-fold, FDR p < 1.32E-16); cellular response to unfolded protein (76.03-fold, FDR p<1.52E-06); and ‘de novo’ protein folding (49.13, FDR p< 2.28E-04). This cluster included small (HSP20-likes) and large heat shock proteins (HSP70, HSP101), their organizing factors (HOP) as well as key regulatory transcription factors HSFA2 (AT2G26150), HSF2BA (AT5G62020), HSFB2B (AT4G11660) and DREB2A (AT5G05410) involved in high light acclimation (see Suppl. Dataset 1, Cluster 2).

Cluster 3 consisted of 123 transcripts with decreased share in the ribosomal and total fractions and was enriched in GO terms for ‘negative regulation of transcription’ (8.91-fold enriched in comparison to the whole genome, FDR p<5.73E-04); ‘response to salicylic acid’ (9.89-fold enriched, FDR p<1.75E-02) and ‘response to oxygen levels’ (10.32-fold enriched, FDR p<5.34E-05). These transcripts included transcription factors WRKY70 (AT3G56400), ERF107 (AT5G61590), as well as growth regulators OFP15 and OFP16 (AT2G36050, AT2G32100, respectively) (see Suppl. Dataset 1, Cluster 2).

Transcripts in Cluster 4 had decreased ribosome association but lacked significant transcriptional changes (Figure 3a). This largest subset among all clusters with 465 transcripts was enriched in multiple GO terms, e.g., ‘response to auxin’ (6.39-fold, FDR p<4.00E-03), ‘tetrapyrrole metabolic process (8.98-fold, FDR p<2.32E-04) and ‘response to far red light’ (4.07-fold, FDR p<8.24E-03), but a predominant response as observed in cluster 1 and 2 was undetectable.

### Plastid translation regulated in response to rapid light shift

While cytosolic translation preferentially recruited photosynthetic and stress-related transcripts in HL, analysis of plastome-encoded genes revealed only the chloroplast-encoded PSBA coding for the PSII subunit D1 as translationally upregulated. Mitochondrial transcripts were missing from the translationally upregulated group. Conversely, 36 plastome- and 16 mitochondriome-encoded DEGs, representing 42.9 % and 34 % of the detected organellar transcripts, displayed a lower ribosome association in cluster 4. The translationally down regulated plastome-encoded transcripts were 50-fold enriched in GO terms for translational processes (30 % in comparison to 0.6 % of genome representation) and 122-fold in ribosome assembly (11 % compared to 0.09 %), but also 100-fold in photosynthetic light reactions (11 % to 0.11 %) represented by YCF3 and YCF4 (ATCG00360, ATCG00520), PSII CP47 reaction center protein (ATCG00680), and NDH-H (ATCG01110). No enrichment in particular GO terms could be detected in the mitochondrial DEGs, but 6 of the 16 DEGs encoded for NADH dehydrogenases.

### Cluster 1 and 2 transcripts eludes global translational repression and MPK6 and AKIN10 regulation

We explored the relationship of transcript abundance (sum of CPM across all samples) and ribosome association (FC ribosome-associated mRNA) of the transcripts for each cluster to query how the mRNA abundance, or availability impacts ribosome association (Figure 3b). Contrarily to the classical translation efficiency (TE) term, this allowed to de-convolute the effects of ribosome loading and unloading from differential transcription (Suppl. Fig. 2c,d). Transcripts of cluster 1 and 4 displayed an inverse correlation between FC of ribosome association and total transcript amount; less abundant transcripts displayed a higher FC in ribosome association (Figure 3b). On the other hand, this relationship was absent in clusters 2 and 3 as FC in ribosome association were independent of total transcript abundance, indicating that ribosome association was dependent on specific transcripts and not purely their abundance.

Given the altered cumulative translational response observed in the ribosome profiles during LL➔HL in *mpk6* and *akin10*, we assessed whether they specifically contribute to cluster 1 and 2 responses. The role of MPK6 and AKIN10 on ribosome association of representative transcripts of cluster 1 and 2 after 10 min LL➔HL was quantified via qPCR of total and ribosome-associated transcripts in *mpk6* and *akin10* (Suppl. Fig. 3). Interestingly, no uniform pattern of deregulation could be observed for this subset of transcripts in the mutants (Suppl. Fig. 3a-d). Querying genotype- and treatment-specific effects via principal component analysis indicated that the treatment and not the genotype accounted for ribosomal association of cluster 1 and 2 representatives (Suppl. Fig. 3e-f). Thus, cluster 3 and 4 represent the cumulative ribosomal regulation as seen in Figure 1c, while cluster 1 and 2 are not represented in the cumulative changes and are unaffected by AKIN10 and MPK6.

Analyzing cluster 1 and 2 transcripts for differential ribosomal association at upstream open reading frames (uORF) revealed only 5 transcripts with shifts to uORFs (Suppl. Table 3), thus no common pattern of changed reading frames of cluster 1 and 2 could be observed. Interestingly, comparing the total RNA-seq read depth with the RPF read depth for enriched maxima of cluster 1 & 2 transcripts revealed enriched ribosomal occupation at certain positions, indicative for ribosome stalling sites (Suppl. Fig. 4).

The increased ribosome association of transcripts of cluster 1 and cluster 2 is in juxtaposition to the global down-regulation of translation after 10 min of LL➔HL. This together with the strong overrepresentation of distinct GO terms in both clusters and the decoupling of transcript abundance and FC in ribosome association, led to the hypothesis that transcript-specific recruitment is taking place. We hypothesized that inherent properties of the mRNA must be co-regulating preferential translation. Therefore, we considered the 5’- and 3’-UTRs of the cluster 1 and 2 transcripts, separately, and combined as translationally upregulated transcripts, for detecting conserved mRNA structures and sequences.

### Translationally regulated transcripts are enriched in RNA motifs in their UTRs

Reorganization of translation requires unloading and reloading of ribosomes with different transcripts. This process is instigated by translation initiation factors acting in concert with mRNA regulatory sequences and/or structures. The 5’-UTR and 3’-UTR possess features that influence the initiation rate, like independent ribosomal entry sites (IRES) [40, 41]. Therefore, the 5’- and 3’-UTRs of the cluster 1 and 2 were screened for conserved motifs enriched in comparison to all TAIR annotated 5’- or 3’-UTRs using the MEME-suite [42]. The search identified 52 motifs in the 5’-UTRS of transcripts in clusters 1 and 2, of which 21 were in common. Three motifs were selected based on their over-representation in cluster 1 and 2 (M1: TCTCCGGAGAA, M2: TTAGGGTT and M3: TCGCCG, Figure 4).

**Figure 4:**
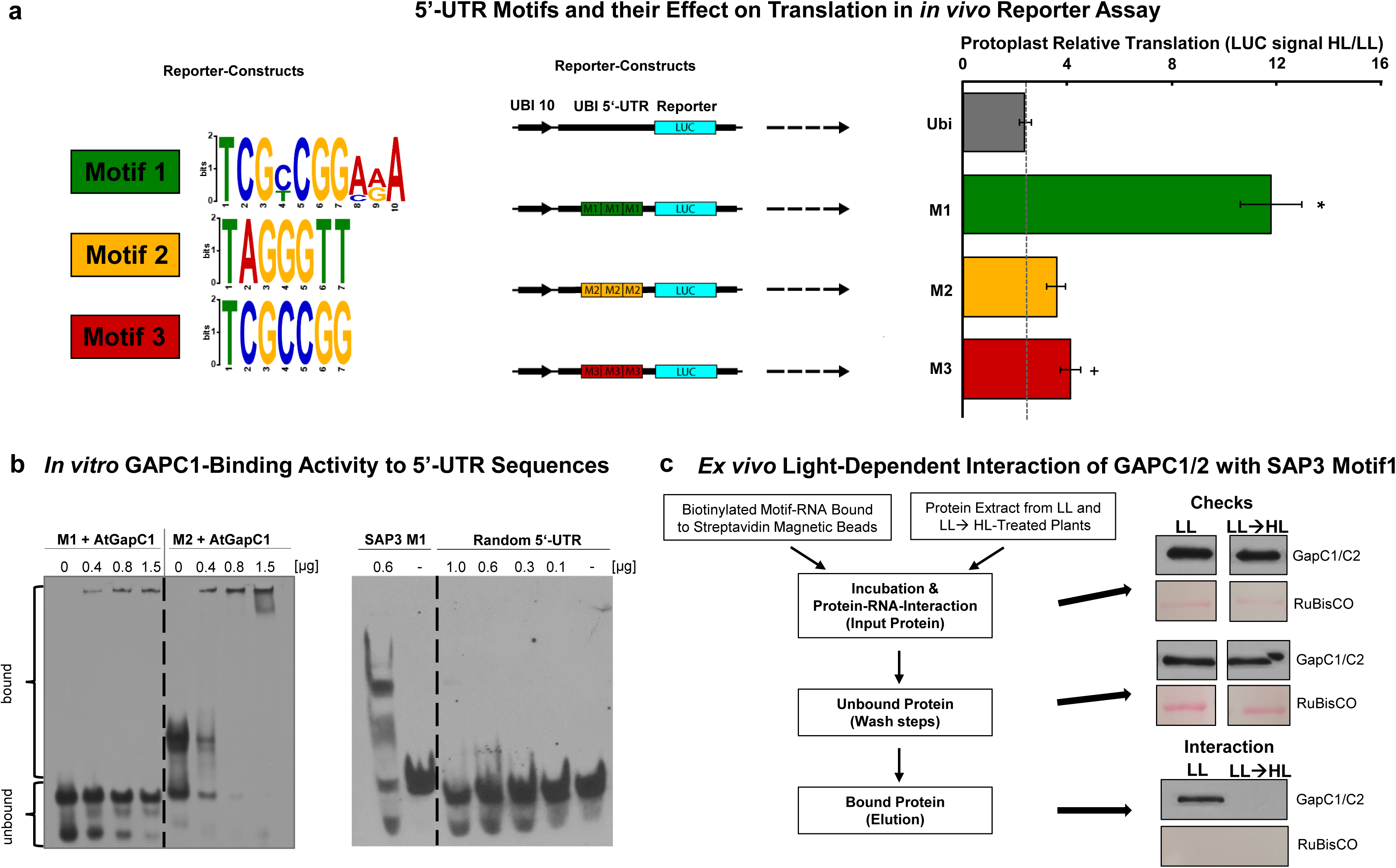
Identification and functional validation of three motifs in the 5’UTR of preferentially ribosome-associated transcripts. These motifs are hypothesized to confer preferential translation after 10 min LL➔HL. (a) In 338 5’-UTRs of the preferentially ribosome-associated transcripts 42 enriched motifs were identified using the MEME algorithm against a randomized 5’-UTR set. Motifs 1 and 3 were present in 32% of those transcripts, and motif 2 in 13% with an additional 8% occurrence in their respective 3’ UTR. The 5’-UTR of *UBI10* was cloned upstream of the luciferase (LUC) reporter under control of the ubiquitin promotor. Triplet motifs were introduced into the 5’-UTR of ubiquitin mRNA. *A. thaliana* mesophyll protoplasts were co-transfected with one of each construct and with the pGL4::RLUC reference construct. Depicted are schematics of constructs used in the reporter assays and relative LUC signal in lysed mesophyll protoplasts treated with HL for 10’ normalized to reference construct. The construct containing M1 showed a strong increase in LUC signal after LL➔HL. Data are means ± SD from two independent experiments with 10 replicates each, * and + indicate significant difference to ubiquitin control, two sided Student’s t-test: p<0.05 and 0.1, respectively. (b) In vitro GAPC1-binding activity to 5’-UTR motifs and random sequences. M1 and M2 were incubated with indicated µg of heterologously expressed AtGAPC1 and subsequently analyzed for RNA electrophoretic mobility-shifts. AtGAPC1 and RNA interaction can be detected by separating at higher apparent size (bound). RNA with M1 and M2 were bound by AtGAPC1 in a dose-dependent manner. This could not be observed for the random RNA oligomer (Random RNA). (c) *Ex vivo* light-dependent interaction of GAPC1/2 with *SAP3* M1. Total protein extract from LL and LL➔HL-treated plants was incubated with biotinylated M1 motif RNA and bound to streptavidin magnetic beads. The input GAPC1/2 protein was detected via SDS PAGE and Western Blot using a GAPC1/2 antibody as well as Ponceau staining of RuBisCO. The magnetic beads were washed from unbound protein and RNA interaction (Wash steps). The bound protein was eluted and a strong GAPC1/2 signal only appeared in the LL protein sample.

Motif M2 was previously identified as stress-responsive 5’-UTR element that confers preferential translation in vitro [43]. M1 and M2 were present 40- and 34-times in the 5’- and 3’-UTRs of cluster 1 and 2 transcripts, accounting for 31% and 26% of transcripts, respectively (Suppl. Fig. 5b). M2 is present in transcripts that are exclusively translationally regulated and not transcriptionally (cluster 1), whereas M1 and M3 are also present in transcriptionally regulated transcripts (cluster 2). Transcripts carrying M1 and M2 had a median FC of relative ribosome association of 2.09 and 1.97, respectively, compared to the median of 2.96 for all cluster 1 transcripts. Transcripts carrying M3 had a higher FC of 3.57. Analysis of motif position in the respective 5’-UTRs of motif-carrying transcripts revealed a preferred position at the 3’-end in longer 5’-UTRs, while short 5’-UTRs showed no positional preference (Suppl. Fig. 5a).

Calculating the secondary structure via RNA Shapes Studio [44] revealed that the motifs do not overlap with any conserved secondary structure (Dot-Bracket annotation, Suppl. Fig. 5a, Suppl. Fig. 6). Heat-responsive transcripts like *HSFB2B* and *HSP20*-like proteins, as well as other stress-dependent transcripts, were overrepresented among the motif-carrying transcripts. Interestingly the 5’-UTRs of the *Stress Associated Protein 3* (*SAP3*) carries all three motifs and that of *SAP2* carries M2 (Suppl. Fig. 5a,b). SAPs belong to the family of zinc finger proteins that contain A20/AN1 zinc-binding domains. They are structurally and functionally conserved among plants [45]. *SAP* transcript amounts increase in response to multiple stresses, and overexpression of specific SAPs in planta enhances biotic or abiotic stress tolerance or photosynthetic capacity [46]. To assess importance of the motifs on an evolutionary scale, we queried their presence in the rice genome. *OsSAP8* and *OsSAP9* were identified as putative homologues of *AtSAP2* and *AtSAP3*, in which we detected minor variants of all three motifs in the corresponding 5’-UTRs (Suppl. Fig. 8a). Additionally, we observed the preferential association of transcripts for *OsSAP8* and *OsSAP9* with polysomes upon L➔H transfer in rice (Suppl. Fig. 8b).

### Enriched 5’ UTR motifs enhance HL-dependent protein synthesis and are targeted by RNA binding proteins

To confirm a regulatory role of the three motifs towards translational regulation, we performed a protoplast-based luciferase assay. Cassettes carrying the M1, M2, or M3 motifs in triplicate repeats were analyzed for translationally regulatory elements and secondary structure and inserted into the *AtUBI10* 5’-UTR (Suppl. Fig. 6) upstream of *luciferin 4-monooxygenase* (*Photinus pyralis*). The verified constructs were transiently co-transfected with pGL4:: *luciferin 2-monooxygenase* (*Renilla reniformis*), as an internal reference, into *A. thaliana* mesophyll protoplasts (Figure 4a). The protoplasts were treated analogously to the whole plant experiments, comparing protoplasts exposed to L➔H-transfer for 10 min to those at LL. The normalized luciferase activity from lysed protoplasts transformed with the ubiquitin 5’-UTR without motifs increased 2.5-fold after L➔H-treatment (Figure 4a). All motifs conveyed a light induced higher signal than the ubiquitin control with M1-construct causing the highest increase of 5-fold, 1.75-fold for M3 and insignificantly increased for M2 (1.52-fold).

We performed RNA-EMSAs to query for potential RBP-binding properties of the motifs. Biotinylated M3-RNA was incubated with leaf protein extracts that were either prepared from plants under LL or exposed to 10 min L➔H. Addition of protein extracts from LL exposed leaves resulted in a pronounced shift of signals in comparison to the biotinylated M3-RNA control (Suppl. Fig. 7a). Addition of unlabeled competitor M3-RNA strongly suppressed the shift. Supplementation with extract from 10 min LL➔HL-treated plants resulted in a signal intensification that again was out-competed by competitor M3-RNA. This result suggests RBP binding capability of M3 with differences in protein signature in extracts from L-light plants and 10 min LL➔HL-plants. To identify the potential RBPs interacting with M1 and M2, an explorative affinity pulldown using total protein extracts from plants under LL and LL➔HL and subsequent SDS-PAGE was performed. The respective enriched bands (Suppl. Fig 7b) were then analyzed via MALDI-TOF. The identified peptides were filtered for peptides matching the respective SDS-PAGE band size and a Mascot-Score of > 60 resulting in GAPC2 and ATP synthase subunit alpha as matches for M1 and M2, respectively.

To confirm this interaction, we performed an RNA-EMSA incubating recombinant GAPC1 with free consensus M1- and M2-RNA (Figure 4b). As demonstrated by a concentration-dependent band shift, GAPC1 could be confirmed as an RBP with binding affinity to consensus M1- and M2-RNA Additionally, GAPC1 did not bind to the motif free 5’-UTR of a transcript that was transcriptionally upregulated during LL➔HL, demonstrating a motif-specific mRNA interaction (Figure 4b). To confirm RNA-RBP interaction with endogenous M1, recombinant GAPC1, GAPC2, or GAPDH isolated from rabbit muscle was incubated with native M1 motif of the *SAP3* 5’-UTR with concentration dependent shifts, which could be suppressed by adding unlabeled competitor RNA (Suppl. Fig. 7c). To further test *ex vivo* interaction in a light dependent context, native M1 motif of the *SAP3* 5’-UTR was used in an affinity pull down with total protein extract form LL and LL➔HL-treated plants, with subsequent immunodetection of GAPC1 and GAPC2 (Figure 4c). Interestingly, a strong GAPC signal could be detected in the affinity pull down from LL total protein extract. In a converse manner, the signal from HL protein extract was weak, with no detectable differences in total GAPC1/C2 content between LL and HL total protein extracts, strongly suggesting a light-dependent interaction of M1 and GAPC1/C2 (Figure 4c).

### Synchronization of translational signaling and nuclear transcription: SAPs interact with CML49 altering nuclear transcription

GO analysis of cluster 2 transcripts revealed an enrichment for stress-responsive transcription factors and co-factors like *MBF1c*, *HSF2A*, and *DREB2A*. These are known to transcriptionally regulate major stress response genes like *HSPs* and *APX2* [47, 48]. Therefore, we hypothesized that rapid translational regulation of cluster 2 transcripts could integrate *de novo* protein synthesis and transcriptional regulation during HL. Next to the afore mentioned well researched transcription factor involved in major stress responses, other targets of translational control were *SAP2* and *SAP3*. With SAP2 fused to the GAL4-DNA-binding domain as bait, six clones were recovered from prey consisting of *A. thaliana* cDNA library fused to the GAL4 activating domain. Five of these clones coded for Calmodulin-like 49 (CML49). The interaction between CML49, and SAP2 and SAP3 was confirmed using the mating-based-Split-Ubiquitin (mbSUS) yeast-two-hybrid system (Figure 5a). Additionally, we performed fluorescence resonance energy transfer (FRET) confirming interactions between SAP2, SAP3, and CML49 with a high FRET efficiency (> 5-fold of negative control) proving their interaction *in vivo* (Figure 5b,c). Interaction of the SAPs with CML49 resulted in trans-localization to the nucleus (Figure 5b). This interaction was also confirmed for OsSAP8/OsSAP9 and OsCML49a/OsCML49b in rice (Suppl. Fig. 9a,b), thus this interaction appears to be a mechanism conserved in flowering plants.

**Figure 5:**
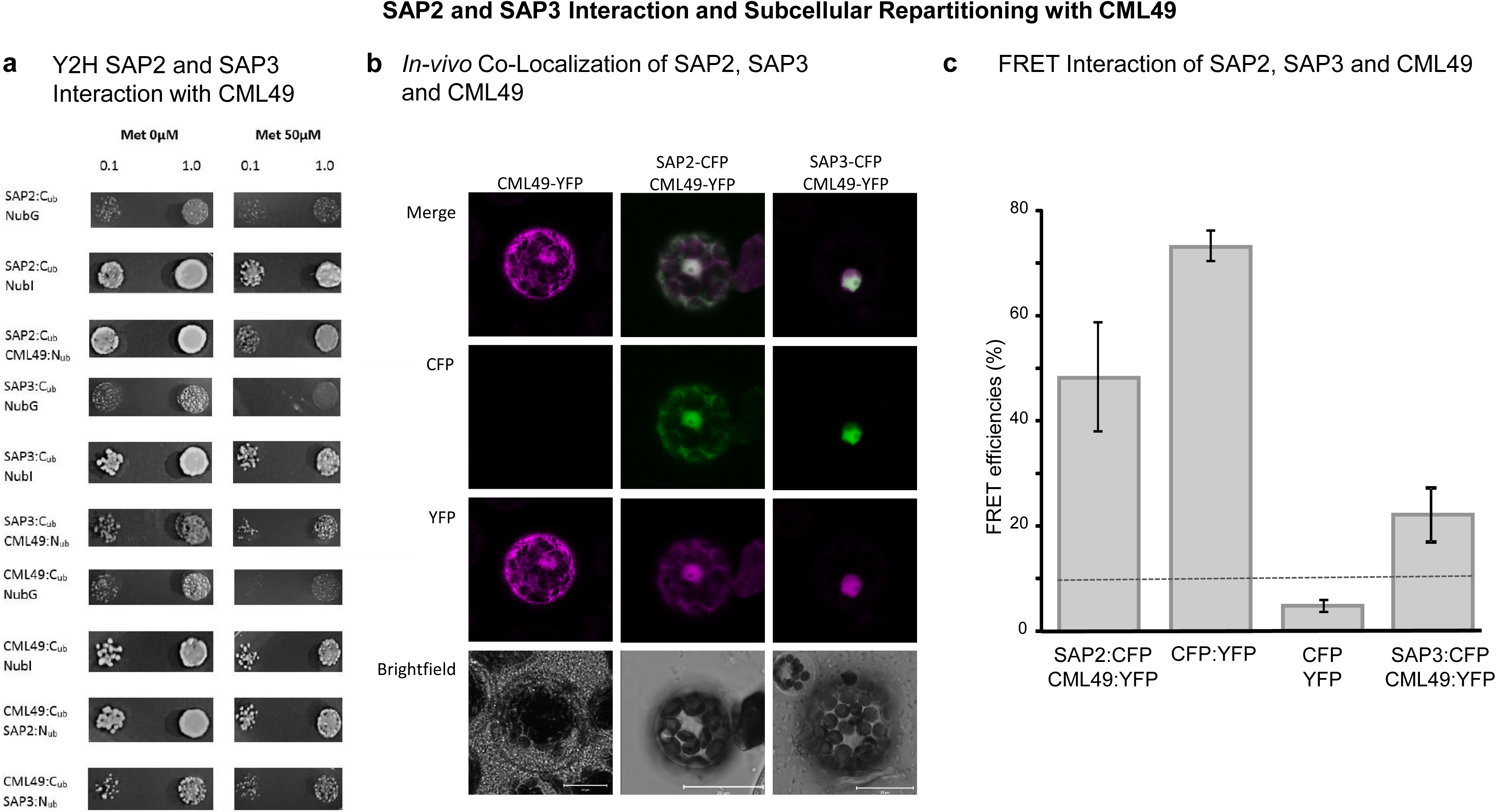
Interaction of SAP2 or SAP3 with CML49 in the split-ubiquitin yeast-two-hybrid system and by Förster resonance energy transfer (FRET) of SAP2:CFP or SAP3:CFP with CML49:YFP in *A. thaliana* protoplasts. (a) Quantification of interaction strength between SAP2 or SAP3 and CML49 using a methionine-dependent split-ubiquitin-assay. Depicted are colonies after 72 h at 30 °C in different ODs (OD = 0.1 and 1.0). Growth was monitored either on Met 0 µM (no methionine, left colonies) or Met 50 µM (50 µM methionine, right colonies) medium plates. Diploid yeast was transformed with NubI- (pNubWtX-gate) as positive control and as negative control with the NubG-vector (pNX35-Dest) as prey (depicted are representative images of n=3 independent experiments). (b) *A. thaliana* mesophyll protoplasts were transfected with SAP2:CFP, SAP3:CFP or CML49:YFP vector DNA. Fluorescence images taken by CLSM for subcellular localization. Following transfection with SAP2:CFP or SAP3:CFP and CML49:YFP constructs, FRET efficiencies between CFP and YFP were quantified. FRET was also determined with CFP:YFP as positive control and 35S::CFP with 35S::YFP as negative control. For each set of transformation more than 20 protoplast were measured twice in n=3 independent transfection experiments.

The preferential association of *SAP2* and *SAP3* with polysomes after LL➔HL-transfer and the interaction of SAP2 and SAP3 with CML49 led us to hypothesize that CML49 and SAP2/SAP3 function as regulatory module in this specific environmental context. This is underlined by the stress-dependent expression pattern for *SAP2* and *SAP3* and the constitutively expressed CML49 within arabidopsis and rice (Suppl. Fig. 10a-f). In addition, the similar response of *SAP2* and *SAP3* in the LL➔HL-response might indicate some functional redundancy. With these two data-guided hypotheses in mind, we hypothesized that the CML49 knockout should have stronger effects on downstream events during H-light acclimation than deletion of SAP3. To investigate the importance of SAP3 and CML49 during HL acclimation, we determined the transcriptional responses of *cml49* and, *sap3* to 60 min LL➔HL (Suppl. Data Set 1). A set of 32 transcripts was more than 3-fold increased or decreased in *sap3* relative to WT, while the corresponding set comprised 214 transcripts in *cml49*.

The data set was examined for transcripts that exhibited stronger downregulation in *cml49* than in *sap3* upon LL➔HL-transfer, since *SAP2* and *SAP3* exhibit similar LL➔HL responses which could be indicative for functional redundancy. This filter revealed 49 transcripts with the expected response pattern (Figure 6a). The predominant GO terms belonged to the categories ‘response to temperature’ and ‘response to light stimulus’, each of which represented 22 % of the genes, followed by ‘response to hydrogen peroxide’ with 11 % and ‘circadian rhythmicity’ with 6 % (Suppl. Fig. 11). Heat shock proteins and one heat shock factor (*HSFA2*) belonged to the group of heat responsive elements, *sigma factor 5* and *Pseudo response regulator 9* (*PRR9*) to the light responsive elements, *Purple acid phosphstase 17* (*PAP17*) and *GDP-L-galactose phosphorylase (VTC2)* to the group of ‘response to hydrogen peroxide’. *Circadian clock associated D 1* (*CCA1*) and *PRR9* represent two genes associated with the circadian clock. The occurrence of transporters for phosphate, sugar and sulphate (*PS3*, *SWEET 13*, *GPT2*, *APR3*) in this list indicates a function of CML49 and SAP3 in regulating metabolite and nutrient distribution. Interestingly, the *Translation elongation factor 1B* (*eEF1B*) was also found in this list possibly pointing to SAP3/CML49-dependent amplification effects in the HL-dependent control of translation.

**Figure 6.**
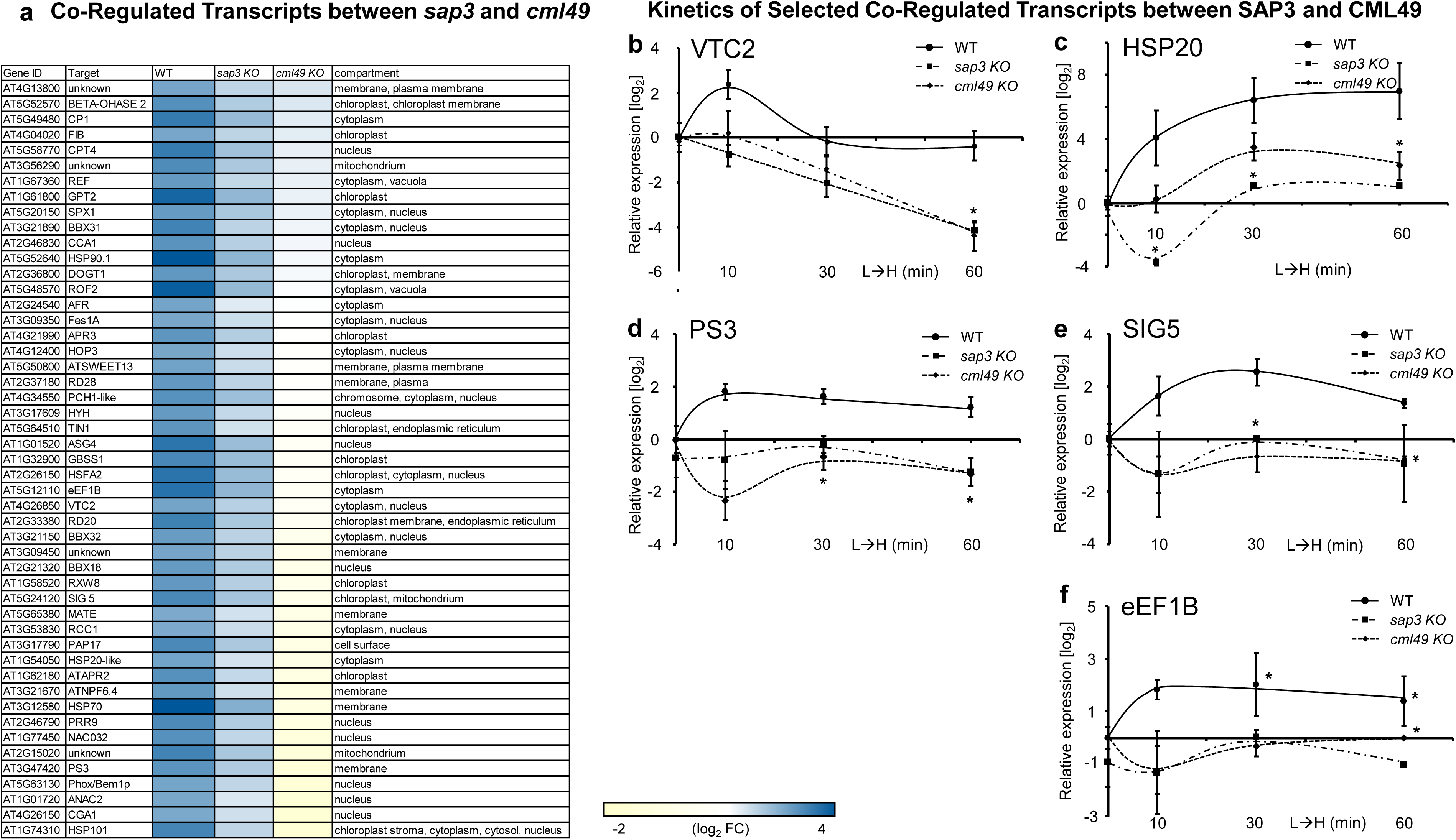
Deregulated HL-response in *sap3* and *cml49* compared to WT. (a) Microarrays were hybridized with cDNA from WT, *cml49* and *sap3* after 60 min of LL➔HL transfer. Assuming a concerted function of SAP2 and SAP3 together with CML49 in the HL response, the data were filtered for co-regulated targets that were stronger deregulated in *cml49* than *sap3* background. The expression level corresponds to the colour intensity with yellow = −2 and blue = 4 (log_2_ fold). (b-f) Validation of transcript responses to HL of putative CML49 and SAP3 targets. VTC2, HSP20-like, PS3, SIG5 and eEF1B were selected based on the transcriptome response, and their transcript levels quantified by qRT-PCR following the LL➔HL-transfer. Data are means ± SD of n=4 independent experiments, asterisks indicate significant difference, two sided Student’s t-test p<0.05.

The next attempt to validate the *SAP3* and *CML49* transcriptional regulation and the kinetics of HL induction (Figure 6b-f) included more detailed analysis of five targets that were selected from the list of co-regulated transcripts. *VTC2* showed a transient four-fold upregulation at 10 min LL➔HL in WT, which dissipated by 30 min. Conversely, *VTC2* was steadily downregulated in *sap3* and *cml49* to four-fold below the L-light level (Figure 6 b-f). *HSP20*-like rapidly increased in WT upon LL➔HL-transfer reaching its maximum of 64-fold induction at 30 min LL➔HL. The mutants showed a slower and lower upregulation than WT with significantly reduced responses at 10 and 30 min LL➔HL in *sap3*, and at 60 min in both mutants (Figure 6b-f). The same trend appeared in WT for *PS3*, *SIG5* and *eEF1B* with a four-fold upregulation after 10 until 60 min of LL➔HL shift. The mutants showed a strong and consistent deregulation compared to WT (Figure 6b-f).

### Common cis-elements in promotor region of co-regulated genes

The 49 co-regulated target genes were screened for common cis-elements in the promotor region in order to find commonalities, such as involved transcription factors required for co-regulation. A total of 9,459 common cis-elements were identified in the 1000-bp upstream of transcriptional start site of the 49 co-regulated genes (Suppl. Table 3). Most abundant common cis-elements were the WBOX, targeting sequences for the WRKY-TFs (2112), followed by sulfur response (SURE, 1536) and MYB and MYB-related cis-element sequences (1368). Stress related cis-elements like the sequences overrepresented in light-induced promotors (SORLIPs) and early responsive to dehydration (ABRE) were also highly abundant with 1278 and 1148, respectively. bZIP cis-elements (C-box, 232), phosphate starvation (PHR1, 224) and salicylic acid inducible (LS7, 173) were less abundant.

### Is there a translational retrograde – anterograde circuit?

After 10 min LL➔HL all plastidial translation was repressed with the sole exception of *PSBA*. This correlated with increased ROS within chloroplasts and subsequent inhibition of plastidial translation [9, 49]. De-repression of plastidial transcription and translation during stress is mitigated via sigma factors like SIG5 which contributes towards transcription of the psbD operon (PSII D2) [50]. Translational regulation of stress responsive transcription factors like HSF2A, DREB2A, and the SAP2/3 interaction with CML49 leads to translation-dependent nucleus synchronization after 10 min LL➔HL. Plastid-targeted proteins like SIG5 and GPT2 and VTC2 were transcriptionally upregulated in CML49- and SAP3-dependent manner but not preferentially translated after 10 min LL➔HL (Figure 6b-f, Suppl. Dataset 1). It is tempting to hypothesize that the SAP-CML49 co-regulated expression of SIG5 contributes to de-repression of chloroplast genes, especially since PSII D2 transcription is 1.4-fold increased after 60 min of LL➔HL.

## Discussion

Regulation of actively translated transcripts described herein adds an entirely new mechanism to retrograde communication in the cell. We identified components of the novel pathway of translation-dependent retrograde signaling (TraDeRS) that bifurcates into direct translation of chloroplast-targeted proteins independently of nuclear transcription (Figure 7 I). In addition, retrograde control of cytosolic translation affects synthesis of a suite of transcription factors that, in turn, promote nuclear transcription (Figure 7 II). Both mechanisms culminate to an anterograde response that enables acclimation. Collectively, the suite of chloroplast-targeted and chloroplast-encoded proteins controlled by TraDeRS includes those involved in protein import and folding, chloroplast transcription, photosynthesis and photoprotection.

**Figure 7:**
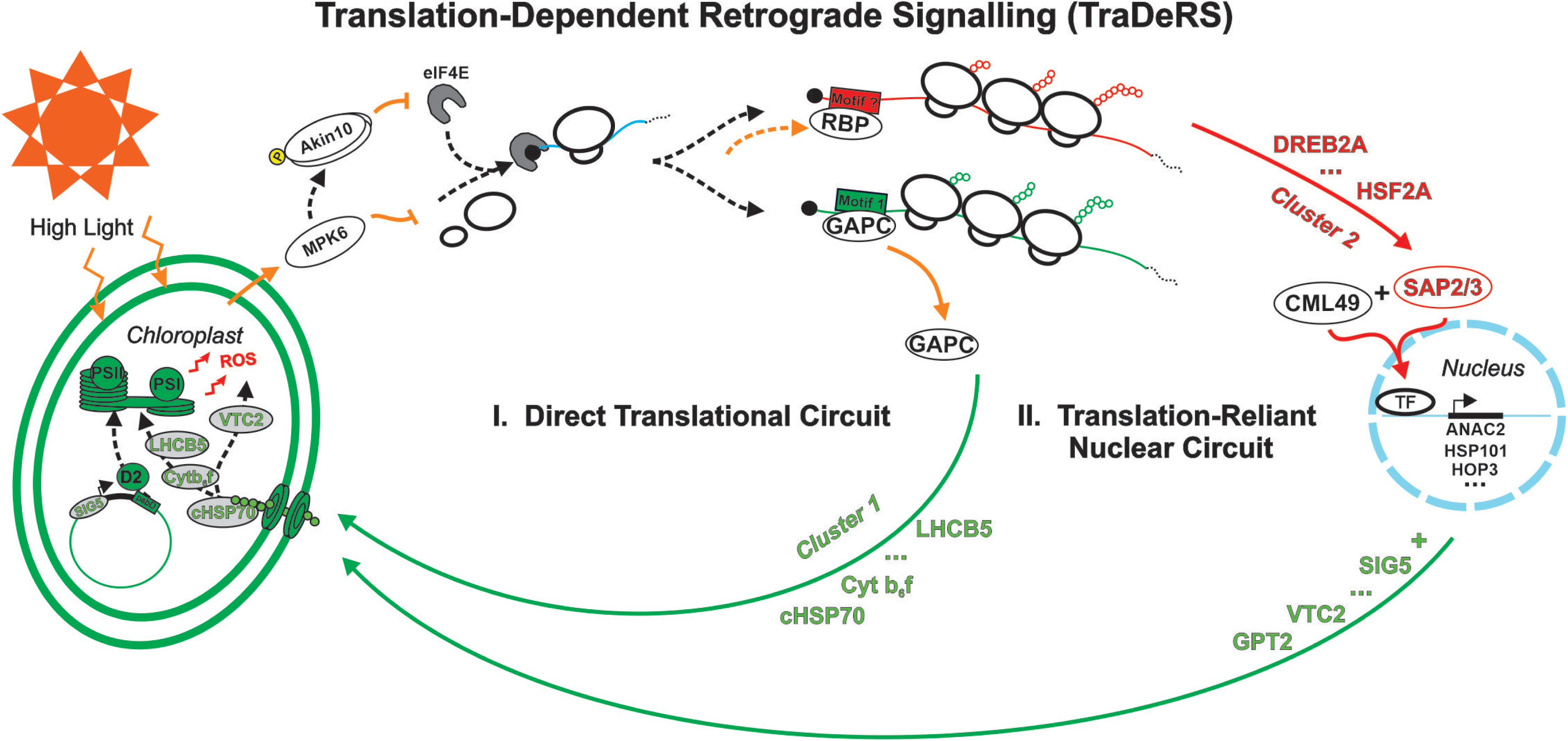
Schematics of translation-dependent retrograde signalling (TraDeRS). Rapid regulation of polysome loading and unloading after 10 min of LL➔HL shift was partially disturbed in *mpk6* and *akin10* mutants indicating retrograde regulation of translation. Regulation involves transcription-independent changes of translational efficiency of majorly chloroplast-targeted proteins involved in photosynthesis as well as chloroplast-associated assembly, repair and import of proteins, thus forming a direct feedback circuit regulated via translation (I. Direct Translational Circuit). The preferential association of transcripts from transcription factors as well as *SAP2* and *SAP3* are part of the translation-reliant synchronization of the nucleus with the changed chloroplast demand (II. Translation-Reliant Nuclear Circuit). SAP2 and SAP3 interact with CML49 and this regulatory module affects transcript abundance of, e.g. *VTC2* and *SIG5*, participating in anterograde signalling, and other major stress response genes like *ANAC2*, *HSP101* and *HOP3*.

### Evidence for translation-dependent retrograde signalling (TraDeRS)

HL stress perturbs and cumulatively represses global translation, while simultaneously increasing translation of stress-specific proteins within minutes of LL➔HL-transfer (Figure 1a-b, Figure 3a). Our work demonstrates that this process is rapidly and reversibly controlled in a light dose-dependent manner (Figure 1a-b). Most other immediate short-term responses to HL are chloroplast-located and post-translational modulated like non-photochemical quenching [51], malate valve stimulation [52], or activation of antioxidant metabolism [53]. In contrast to the chloroplast-localized processes, TraDeRS takes place in the cytosol and potentially integrates multiple cues for changes in *de novo* protein synthesis. Previously, we demonstrated that rapid transcriptional responses in the nucleus affect a network of transcription factors within 20 s after the LL➔HL-transfer [9, 54]. Only later peaks the H_2_O_2_ burst at 30 min after the LL➔HL. The decreased polysome association of transcripts displayed a reciprocal behavior with the H_2_O_2_ burst after the LL➔HL shift. The maximal H_2_O_2_ accumulation after 30 min is followed by a return to basal levels after 60 min [9]. Parallel to the decreasing H_2_O_2_-content, general ribosome association of mRNA recovered after one hour of HL (Figure 1b and as reported [43]).

### Retrograde-dependent translational control involves MPK6 and AKIN10

Ribosome unloading and preferential loading of transcripts are decisive steps in translational control in the cycle of translation initiation, elongation, and termination [55]. The question is how does HL perceived in the chloroplast modulate translation? Protein phosphorylation, thiol redox-related changes and acetylation are reported post-translational modifications (PTM) that regulate translation initiation, elongation and termination factors [56]. Formation of the translational pre-initiation complex (PIC) is inhibited by the GCN2 [57], and its activation has been shown in to be linked to MAPK signaling pathways [58]. Consequently, our results showed that mutants lacking MPK6 exhibit an altered ribosomal profile. Although the global association of transcripts to ribosomes were altered in *mpk6* (Figure 2) hinting at altered translation initiation, none of the marker transcripts from the increased translational cluster were disturbed in their preferential ribosome association upon LL➔HL-transfer (Suppl. Fig. 3a-d). This is indicative of a stress-dependent separation of global inhibition of translation initiation via MPK6 and the increased ribosome association of specific transcripts through a distinct mechanism.

The second rate-limiting mechanism of translation initiation is via eIF4F complex formation and subsequent mRNA binding regulated via 4EBPs. In mammals 4EBPs are activated by the mammalian Target of rapamycin (mTOR) kinase [59] mediating phosphorylation of the eukaryotic Initiation factor 4E (eIF4E), disabling interaction with initiation factor eIF4G, and thus limiting eIF4F formation [60]. Homologous 4eBPs being target of TOR signaling pathways in plants could not be identified yet. However, phosphorylation of the Translation initiation factors eIF4E and eIFiso4E via SnRK1, an interactor of TOR, inhibited translation in a plant system [61]. Herein we show that the lack of AKIN10, a subunit of SnRK1, altered the ribosomal profile, under control and stress, and interfered with the increased ribosome association of the marker transcript *SAP2* (Suppl Fig. 3a). A future question is what activates this pathway? ROS or redox-mediated regulation are candidates, as are GCN2 and TOR-signaling via AKIN10 and MPK6, and DHAP. We previously reported that TPT2-mediated export of DHAP from the chloroplast was required for rapid phosphorylation of MPK6 after 10 min LL➔HL-transfer. In summary, the results show that both MPK6 and AKIN10 are required for stress-dependent translational control in a retrograde signaling context, most likely via a phosphorylation-dependent pathway. Importantly, the increased ribosome association of specific transcripts appears to occur independently of this pathway.

### Transcripts affected by bifurcating translation-dependent signaling pathways

The set of preferentially loaded transcripts could be grouped into purely translationally regulated and both, translation- and transcription-regulated transcripts, with each group representing distinct pathways of the bifurcated acclimation responses. Transcripts related to photosynthesis, heat stress and H_2_O_2_ dominated the set of preferentially polysome-loaded and motif-carrying transcripts (Suppl. Dataset 1, cluster2). Photosynthesis and chloroplast repair-related transcripts were highly enriched among the purely translationally regulated transcripts (Suppl. Dataset 1, Cluster1). The 119 transcripts in the translation-only regulatory cluster were 5-fold enriched in chloroplast-localized proteins according to SUBA4 [39]. This set includes transcripts encoding proteins critical for assembly, function, and maintenance of the chloroplast. Minor and major Light harvesting proteins, Cytochrome b_6_f (*AT2G26500*) and NADP oxidoreductase (*AT1G06690*) as well as one small subunit of RuBisCO (AT5G38420). Additionally, PSI assembly factor PYG7/YCF37 (*AT2G23670*) and import chaperones like HSC70-1 (*AT5G02500*) and chloroplast vesiculation factors (*AT2G25625*). Our results strongly suggest the existence of transcription-independent regulation of the photosystem repair mechanism that is under rapid translation-dependent control, forming a direct retrograde circuit (Figure 7, I. Translational Circuit).

Complementary to photosystem repair, translational control of nuclear transcription factors (HSFs, DREBs) synchronizes the gene expression with the cellular demand for subsequent stress acclimation. The identified HSF2A, DREB2A and MBF1c play major roles in transcriptional control of HL responses [48, 62]. Interestingly, transcripts for these translation factors are already present under LL but not preferentially translated (Suppl. DataSet 1). Preferential translation after stress perception might serve as first activator of these transcription factors that are known to be up-regulated by oxidative stresses, such as HL.

### Regulatory elements in transcript UTRs controlling light-dependent ribosome association

Transcript-specific translation impacts *de novo* synthesis of distinct protein classes and must be inherently based on regulatory RNA sequence or structural properties. To identify conserved sequences that might serve as such regulatory property, we conducted *de novo* motif detection in UTRs of translationally regulated transcripts (cluster 1 and 2) revealing 42 motifs. We could demonstrate *in vivo* impacts of three identified motifs during HL-stress on translation and provide evidence suggesting that they serve as RBP binding sites for GAPC1/GAPC2 (Figure 4, Suppl. Fig. 7a-c). Indeed, RNA-EMSA shifts with leaf extracts from LL and LL➔HL leaves indicate that M1 acts as an mRNP binding site (Figure 4c). Conditional RBP-binding and stimulation of translation could be shown for motifs M1>>M3>M2 in the arabidopsis protoplast assay (Figure 4a). The motif M2 (TAGGGTTT) has previously been identified in the 5’-UTRs of a set of preferentially polysome-associated transcripts in young seedlings treated with light for 30 min. Translation of transcripts encoding reporters carrying M2 *in vitro* occurred at higher rates than constructs lacking the sequence [43]. The motif was also overrepresented in preferentially polysome-associated transcripts under hypoxic stress [63], suggesting a role for motif M2 under various stresses. However, prior to this study its mode of action had not been reported.

The motif-binding RBPs were investigated by RNA-based affinity chromatography and mass-spectrometry. This identified ATP synthase subunit alpha and GAPC2 as candidate RBPs for M1 and M2, respectively. The other identified peptides were either not matching in predicted protein size, or had a too low Mascot-Score and were excluded as background or unspecific binding. GAPDH functions as non-canonical RBP in animal systems [64, 65] and this role was only recently discussed in plants as well [66]. We validated the GAPDH / motif interaction using the native M1 in the *SAP3* 5’-UTR via *ex-vivo* RNA-affinity chromatography, and subsequent immunodetection, and *in vitro* by RNA-EMSA. The interaction was strong with LL protein extracts and almost lost in extracts from LL➔HL shifted leaves, pointing towards a light-dependent dissociation of GAPC2 (Figure 4c). Therefore, this dissociation upon LL➔HL shift appears to release repression of ribosome association, for transcripts harboring the motifs. As protein amounts of GAPDH (C1/C2) were unaltered by LL➔HL treated plants, conditional RNA binding could be mediated by PTM or the availability of co-factors or substrates. This may link translational control by RBPs to the cellular metabolic state, thereby enabling the cell to rapidly adjust protein synthesis according to metabolic requirements.

### *SAP2* and *SAP3* contain motifs 1 and 2 and their proteins interact with CML49

Motifs M1 and M2 are both present in the 5’-UTRs of *SAP2* and *SAP3* from arabidopsis and also in their homologues in the monocot rice. The transcripts in rice showed the same polysomal association during L➔H as in arabidopsis. Interestingly, re-analyzing a dataset of arabidopsis seedlings re-illuminated after darkness for 10 min [26] for translationally regulated transcripts in the same manner as performed for this study, revealed *SAP2* as one of only five transcripts overlapping with the purely translationally regulated subgroup (cluster1, Suppl. Dataset 1).

The SAPs belong to a zinc-finger multigene family with a highly conserved structure, leading to similarity among several plant species [45, 46]. Despite little molecular knowledge, two regulatory roles have been attributed to SAPs, namely ubiquitin ligase activity and redox sensor activity regulating gene expression under abiotic stress [46, 67–69]. Previous work demonstrated that SAP12 undergoes reversible redox-dependent conformational changes induced during oxidative stress [67]. Given the high level of sequence conservation in the SAP-protein family we analyzed SAP2 and SAP3 for disulfide bonds with the DiANNA [70] tool suit. The analysis revealed a predicted disulfide bond in the AN1 domains of SAP2 and SAP3 at Cys117 and Cys135, respectively. SAP2 had one additional disulfide bond predicted between Cys8 and Cys22. Thus, SAP2 and SAP3 might also function as redox sensors under HL. Furthermore, SAP5 interacts with MBF1c to regulate acclimatory responses, during heat stress via interaction with HSF and DREB transcription factors, ultimately influencing transcription [71]. Given the structural similarity between SAP2, SAP3 and SAP5, the interaction of SAPs with MBF1c might co-regulate transcript specificity while interaction between HSFs, DREBs and MBF1c synchronize transcriptional demand, but this remains subject to further study. Nonetheless, as a consequence of retrograde signaling the abundance of SAP2 and SAP3 is increased and, in parallel, promotes their re-location to the nucleus to promote transcription.

SAP2 and SAP3 interact with the Calmodulin-like-protein-49 (CML49). Several CMLs display an electrophoretic mobility shift in the presence of Ca^2+^, indicating that they act as Ca^2+^ sensors [72, 73]. Most CMLs lack known functional domains that impede the prediction of biochemical functions. Nonetheless, CMLs are important in signaling processes and are activated under stress conditions like heat, cold, drought, and heavy metal accumulation [74, 75]. The Ca^2+^-dependent binding of CaM7 to a Z-/G-box is involved in regulating expression of light-responsive genes [76]. H_2_O_2_ triggers an increase in cytosolic Ca^2+^ by activating Ca^2+^ channels [77]. Ca^2+^-liganded CaM activates MPK8 which suppresses ROS accumulation via transcriptional control of *RBOHD* [78]. *CML49* occurs in almost all tissues and was not transcriptionally or translationally induced after 1 h of LL➔HL indicating that *CML49* functions as constitutive signal transmitter. Interestingly, the rice homologues of *SAP2* and *SAP3* (*OsSAP8* and *OsSAP9*) also interact with *CML49* homologues indicating a conserved regulatory mechanism potentially integrating ROS cues with Ca^2+^ signaling (Suppl. Fig. 8). But this discussed function needs further experimental confirmation.

### SAP3 and CML49 interaction co-regulates light-induced transcription

Both *sap3* and *cml49* knockouts display smaller sets of light-induced transcripts compared to WT. We propose that CML49 acts as general interactor with both SAPs, while each SAP acts as a stress-specific partner. First, CML49 is constitutively expressed while *SAP2* and *SAP3* are preferentially translated under HL. Newly translated SAPs can then interact with available CML49 resulting in relocation of the complex to the nucleus. Second, of the 215 transcripts downregulated in *cml49* plants after 60 min of LL➔HL transfer a stress subset of 32 are downregulated in *sap3*. Thus, the stress-responsive protein, SAP3, is providing specificity to what would be otherwise a broader CML49 response (Figure 6a). Cumulatively, the preferential translation of the SAPs and their interaction with CML49, but also other TFs like DREB2A and HSF2A, form a translation-reliant signal pathway affecting nuclear transcription.

This translation-reliant nuclear circuit regulates two major clusters of genes. As noted above, the majority of the transcripts regulated by the proposed CML49-SAP3 module code for stress responsive transcripts. A subset which includes cytosolic *HSP70*, *HOP3*, *ROF2*, *HSFA2*, *HSP90.1*, *HSP20-like* that increase after 1 h of LL➔HL shift (Figure 6 a). Interestingly, *HSP90.1* and *ROF1* interaction is needed for HSFA2-dependent heat memory and also regulates *short HSPs* (*sHSPs*) [79]. HSFs have also been shown to translocate to the nucleus in a redox-regulated manner [80]. Taken together, we propose that this mechanism forms a feed-forward loop that can amplify the transcriptional responses over time, the translation-reliant retrograde stress signaling pathway (Figure 7, II. Translation-Reliant Nuclear Circuit).

A second subset of nuclear-encoded transcripts co-regulated by CML49 and SAP3 takes part in an anterograde signaling pathway that collectively could contribute to chloroplast acclimation. Among the transcripts were *VTC2*, encoding GDP-L-galactose phosphorylase, which catalyzes the rate limiting enzymatic step in ascorbate synthesis [81]. With respect to anterograde signaling, plastidic *Sigma factor 5* (*SIG5*) is utilized by chloroplast-encoded RNA-polymerase to tran scribe the chloroplast-encoded *psbD* gene, encoding the D2 protein of the reaction center of photosystem II [6], that is induced by HL [82]. *SIG5* and *D2* transcripts were increased in response to the LL➔HL, and *SIG5* was reduced in the *cml49* and *sap3* knockouts, providing evidence for an anterograde circuit potentially affecting chloroplast transcription at later time-points in response to translational-dependent retrograde signaling.

## Conclusion

This work reveals a translation-dependent retrograde signaling pathway and identifies major participants across key stages that regulate rapid acclimatory responses to LL➔HL transfer. The translatome reorganization initiated by LL➔HL involves phosphorylation cascades involving MPK6 and AKIN10, which inhibit cumulatively global translation while simultaneously preferential ribosome association occurs for specific functionally distinct transcripts by a different mechanism. This may serve as a very efficient local control mechanism (i.e. circumventing the nucleus) and is possibly faster than transcriptional regulation for synthesizing proteins required for chloroplast acclimation. The preferentially loaded transcripts directly code for repair and photosynthesis-related proteins forming a direct translational retrograde circuit (Figure 7). Preferential ribosome association is facilitated by conserved motifs in their 5’-UTR acting as RBP binding sites. The motifs, together with GAPC2, function in the direct circuit in a light-dependent manner. A translation-reliant nuclear circuit regulates, next to well-studied stress related transcription factors, the translation of SAP2 and SAP3 in response to LL➔HL together with their interaction with the Ca^2+^ sensor CML49 to promote accumulation of stress-responsive transcripts and anterograde signaling via SIG5, which functions in plastid transcription. The elucidated multiple components of a bifurcating translation-dependent retrograde signaling pathway allows for rapid tuning of protein synthesis to account for the demand of the photosynthesizing cell.

## Materials and Methods

### Plant growth and treatments

*A. thaliana* wild-type (ecotype Columbia) and mutants (*cml49-1* (SALK_090873), *cml49-2* (SALK_035905), *sap3-2 (SALK_022340), sap3-3 (SALK_136520C)*, *sap2-1* (SALK_008677), *sap2-2* (SAIL_1182_C11), *mpk6-3* [83], and *akin 10* (GABI_579E09) were grown under short day conditions (10 h light with 80-100 µmol photons m^−2^ s^−1^ (NL), and 21°C/ 14 h dark, and 18°C, 50 % relative humidity) for three weeks prior to transfer to 8 µmol photons m^−2^ s^−1^ (LL) for 10 days. Plants were transferred to 800 µmol photons m^−2^ s^−1^ (HL) for time periods as indicated, while control plants remained in LL. Whole rosettes were harvested under control and HL conditions in the midday point of the light period to exclude circadian effects and immediately frozen in liquid nitrogen and stored at −80°C.

### Polysome profiling and sucrose cushion for polysome isolation

Total ribosomes and ribosome-bound mRNA were extracted for polysome profiling and quality check via TEM after separation by sucrose gradient centrifugation as described [31]. Polysomal profiles were blank gradient normalized and the peaks corresponding to 60S, 80S and polysomes integrated before normalization to WT values. Significant differences were calculated via two sided Students’s t-test p≤0.05. Sucrose cushion centrifugation was performed prior to foot print analysis. 3 g of pulverized plant material was used with 6 mL polysomal extraction buffer (PEB; 0.2 M Tris, pH 9.0, 0.2 M KCl, 25 mM EGTA, 35 mM MgCl, 1 % Brij, 1 % Triton X-100, 1 % Tween 20, 1 % Igepal CA630, 1% sodium deoxycholate (DOC), 1 % polyethylene-10-tridecylether (PTE), 5 mM DTT, 1 mM PMSF, 100 µg/mL cycloheximide, 100 µg/mL chloramphenicol, 100 µg/mL lincomycin). The suspension was passed twice through Miracloth^TM^ and subsequently cleared by centrifugation. Extracts were loaded on a sucrose cushion (0.2 M Tris, pH 9.0, 0.2 M KCl, 0.025 M EGTA, 0.035 M MgCl, 1.75 M sucrose, 5 mM DTT, 50 µg/mL cycloheximide, 50 µg/mL chloramphenicol, 50 µg/mL lincomycin) and centrifuged for 18 h at 100,000 *x*g in a Beckmann SW71Ti rotor. The sediment was suspended in 100 µL RNase digestion buffer (20 mM Tris-HCl, pH 8.0, 140 mM KCl, 35 mM MgCl_2_, 50 µg/µL cycloheximide, 50 µg/µL chloramphenicol, 50 µg/µL lincomycin) and used immediately for RNase If digestion.

### Transmission Electron Microscopy (TEM) Imaging

TEM images were generated at the Centre for Advanced Microscopy at ANU using the JEM F200 (JEOL) by using 2 µL of undiluted sucrose gradient from fractions representing monosomal or polysomal fractions and negative staining as described in Booth et. al. [84].

### RNase If digestion

RNase If digest was performed with 2,000 relative absorbance units OD_260_ of the in RNase digestion buffer suspended polysomes by adding 50 U of RNase If (New England Biolabs) in 250 µL total volume by incubation at room temperature with constant rotation for 1 h. In parallel same amounts of polysomes were treated with 5 µL SUPERase RNase inhibitor (Thermo) and kept on ice as control sample. The digestion was stopped by adding 5 µL SUPERase RNase inhibitor and the RNA precipitated by adding 1 volume of isopropanol, 300 mM NaOAc pH 5.2 and 20 µg glycogen at −80 over-night. The supernatant was discarded after centrifugation at 16,000 *x*g at 4 °C for 30 min and the sediment was washed twice with 75 % ethanol and resuspended in RNA loading buffer for size exclusion electrophoresis.

### Library sequencing and analysis of mRNAs

Total RNA was isolated using Tri Reagent (Sigma-Aldrich) from extracts suspended in polysomal extraction buffer (see sucrose cushion for polysome isolation, above). Briefly, 500 µL of extract was combined with 1 mL of TriReagent, followed twice by extraction of the organic phase with 200 µL chloroform. RNA was precipitated with equal volume 100% isopropanol and incubated overnight at −20°C. The RNA was recovered by centrifugation and washed with 75% ethanol before resuspension in water. RNA quality was assessed using a LabChip GXII (Perkin-Elmer). Total RNA was DNase-treated using TURBO DNase (Invitrogen) following the manufacturer’s protocol. Preparation of total RNAseq libraries was carried out using Illumina TruSeq Stranded Total RNA with Ribo-Zero plant, scaled at half reaction volumes.

Libraries of ribosome-protected fragments (RPFs) were prepared from three biological replicates per time-point and sample following RNase If digestion (above) according to Juntawong et al. (2014) with minor modifications. RNA linker, rRNA subtraction, and PCR oligonucleotides were ordered from Integrated DNA Technologies (IDT) and are listed in Suppl. Table 2. Oligonucleotides for rRNA subtraction were adapted from those listed in [36] and [85]. Additionally, rRNA1c, rRNA1 and rRNA2 depletion primers were custom designed targeting the highest abundant rRNA detected during method establishment sequencing.

Total RNA and RPF sequencing libraries were quantified using the Qubit fluorometer (Invitrogen) and size distribution was assessed using the LabChip GXII. Samples were pooled and sequenced (75 bp single end) on the NextSeq500 at the Biomolecular Resource Facility (BRF) at the Australian National University.

Quality control of sequencing data was performed with FastQC (v.0.11.5). Total RNA data had adapters removed using scythe (v.0.991) with a prior of 0.01, and quality trimmed with sickle (v.1.33) with a quality threshold of 20 and a minimum trimmed length of 20. Mapping to TAIR10 was performed with subjunc (v.1.4.5), reporting only reads with a single unambiguous mapping location [86]. RPF data was trimmed for adapter contamination and read quality using Cutadapt (v.1.9.1) with a minimum trimmed length of 20 bp and error rate of 0.1 [87]. Trimmed reads were mapped to TAIR10 using Bowtie2 (v2.2.9) requiring end-to-end alignment and a substring length of 10 [88]. Total and RPF-mapped reads were summarised using featureCounts [89]. Counts data was converted to reads per kilobase million (RPKM), and counts per million reads mapped (CPM) in R using edgeR [90, 91]. Only transcripts with counts of RPKM above 7 for all three replicates were considered for subsequent analysis.

### DEG definition and transcript clustering

Cut-offs for assignment as Differentially Expressed Genes (DEGs) were chosen in a data-driven process, according to upper and lower quantiles from a random distribution analysis of all regulated polysome-associated and total transcripts (total mRNA: 1.5 and 0.75 and polysomal: 1.1 and 0.75; 0.05 error margin). Clusters were derived for transcripts significantly changed up and down in both polysomal and total RNA (cluster 1 and 4). Cluster 2 and 3 were selected by filtering for only significantly up- and down-regulated in the polysomal set and not significantly changed in total RNA. Other transcripts were not assigned to clusters, such as 33 DEGs that were changed in total mRNA but not in the ribosomal fraction and termed as buffered transcripts.

### Whole genome ATH1 microarray

Total RNA was isolated from 1 h LL➔HL-treated *sap3*, *cml49* and WT plants using RNeasy mini Kit (Qiagen). Controls were kept in LL. The hybridization of the Affymetrix ATH1 genome Array was commissioned to KFB (Regensburg, Germany). Each condition was hybridized three times from independent experiments for *sap3* and WT and once for *cml49*. Microarray data were analysed with ROBIN (MPI Golm, Potsdam, Germany) and normalized with RMA algorithm [92]. Transcripts were considered as differentially regulated if the FC was >2 and the adjusted *p*-value <0.05 [93, 94].

### Transcript profiling

RNA isolation and cDNA synthesis were performed as in [95]. Quantitative real-time PCR (qPCR) of total and polysomal transcripts was conducted as described in [96] with the difference of using KAPA SYBR^®^ FAST qPCR Kit (Kapa Biosystems, South Africa). *ACTIN2* (*AT3G18780*) and *TUBULIN5* (*AT1G20010*) were used as reference genes.

### RNA electrophoretic mobility-shift assay (RNA-EMSA)

5’-biotinylated RNA and unmodified RNA of Motif 3 (5’-AAAUCGCCGGAG-3’ 5’-Bio-AAAUCGCCGGAG-3’) were synthesized by Eurofins (Hamburg, Germany) and used for LightShift™ Chemiluminescent RNA EMSA Kit (Thermo Scientific). Total protein was extracted from LL and 10 min LL➔HL-treated plants using extraction buffer (25 mM Tris, pH 7.8, 75 mM NaCl, 15 mM EGTA, 15 mM glycerophosphate, 15 mM 4-nitrophenyl pyrophosphate, 10 mM MgCl_2_, 1 mM DTT, 1 mM NaF, 0.5 mM NA_3_VO_4_, 0.5 mM PMSF, 10 µg/mL leupeptin, 10 µg/mL aprotinin, 0.1 % Tween-20). Prior to separation by non-denaturing polyacrylamide gel electrophoresis, RNAs were incubated with extracts at room temperature for 30 min. Biotinylated RNA was detected by chemiluminescence with streptavidin-HRP. For targeted EMSA AtGAPC1 and ArGAPC2 were expressed in *E.coli* BL21 (DE3) pLysS and purified as described in [97] with slight modifications. Buffers were not supplemented with NAD^+^ and purified protein was stored in 10 mM NaPi, pH 7.4. AtGAPC1, AtGAPC2 and GAPDH from rabbit muscle (SIGMA Aldrich) were used for mobility shift assays with 5’-biotinylated M1-RNA (5’-BioTEG-AUAUAUUCUCCGGAGAA-3’), 5’ biotinylated M2-RNA (5’-BioTEG-AUAUAUGAGAUUAGGGUUU-3’), 5’-biotinylated SAP3 M1 (5’-Bio-ACCAGAUUUUUUUCGCCGGAGUUUGUUUUGAUCCA-3’), and random 5’UTR (*AT5G16470*) with mutated M2-sequence (5’-Bio-UUAGCGUUUUAAU*GUUGAGUGAGUA*AGUUAAGGGGUU-3’).

### Reporter gene construct and transient protoplast assay

For the reporter gene construct assay solely *UBI* 10 5’-UTR as control (CCATGGCTTGATCACGGTAGAGAGAATTG–AGAGAAAGTTTTTAAGATTTTGAGAAATTGAAATTCTCG) or *UBI* 10 5’-UTR with motifs (M1: CCATGGCTTGATCACGGTAGAGAGAATTGAGAGAAAGTTTTTTTC–TCCGGAGATTCTCCGGAGATTCTCCGGAGAAAGATTTTGAGAAATTGAAATTCTCGAG; M2: CCATGGCTTGATCACGGTAGAGAGAATTGAGAGAAAGTTTTTAAACCC–TAATCTCAAACCCTAATCTCAAACCCTAATCTCAAGATTTTGAGAAATTGAAATTCTC GAG; M3: CCATGGCTTGATCACGGTAGAGAGAATTGAGAGAAAGTTTTTCTCCGG–CGACTCCGGCGACTCCGGCGAAAGATTTTGAGAAATTGAAATTCTCGAGA) were cloned into MS129 luciferase reporter [98], and 5 μg plasmid DNA was co-transfected with same amount of low-rate constitutively expressed pRL-TK *Renilla* luciferase vector (Promega) into mesophyll protoplasts isolated from *A. thaliana* leaves by the downsized polyethylene glycol-mediated transfection method (Seidel et al., 2004). After 12 h of acclimation protoplast were transferred to LL and half of the transfected protoplasts were transferred to HL for 10 min and subsequently flash frozen in N_2 liq_ for cell extracts. Luciferase activity of cell extracts was measured with Sirius L Tube Luminometer (Berthold) and the luciferase assay system (Promega).

### RBP pulldown and identification

Affinity pulldown with motif-carrying biotinylated RNA for detection and identification of potential protein-RNA interactors was conducted as described in [99] with minor differences. In short: 500 µg (or 250 µg for pull down followed by immunodetection of cytosolic GAPDH) of total protein from LL and LL➔HL-treated plants were incubated for 30 min at RT with 1 µg biotinylated M1- or M2-RNA (3 µg for 5’-Bio-*SAP3*-M1), 1 µL aprotinin (25 µg/µL), 1 µL leupeptin (25 µg/µL), 2 µL RNasin (Promega), 250 µL TENT buffer (0.02 M Tris-HCl, pH 8.0), 0.002 M EDTA (pH 8.0), 0.5 M NaCl, 1 % (v/v) Triton X-100) and adjusted to 500 µL with TENT buffer. 50 µL Dynabeads (Thermo Fisher) were added and incubated under slow shaking for 30 min at RT. After precipitation and three washing steps with TENT buffer, Dynabeads were dissolved in 1x protein loading buffer and incubated for 10 min at 95 °C, precipitated and the supernatant loaded on SDS-PAGE for separation and band identification via mass-spectrometry or immunodetection of GAPC1/C2 after blotting the gel-separated proteins on nitrocellulose membrane by semi dry electro blotting technique.

### Promotor and uORF analysis

The +50 to −1000-bp region upstream of the ATG start codon and +500 downstream of the transcription stop site of the CML49 and SAP3 co-regulated genes were analysed for common *cis*-elements using PlantPAN2.0 [100]. For uORF analysis the web tool uPEPperoni was used [101] and the eFP Browser 2.0 for general expression pattern [102].

### mbSUS-analysis

Vector construction of the mating-based Split-Ubiquitin-System analysis was performed with the Gateway® cloning system (InvitrogenTM). The coding sequences of SAP2, SAP3 and CML49 were fused to attB recombination sites and cloned into the pDONRTM221-vector by a BP-reaction. Subsequently, each entry clone was used in a LR-reaction with the destination vectors pX-NubWTgate (Nub) and pMetYCgate (Cub). Controls were the transformed diploid yeast with NubI- (pNubWtX-gate, positive control) and the NubG-vector (pNX35-Dest, negative control) as prey. The resulting vectors were transformed via lithium acetate transformation into *Saccharomyces cerevisiae* strains THY.AP5 and THY.AP4, respectively [103]. Yeast cells were grown in 20 mL selective Nub/Cub medium at 30°C/140 rpm for 5-7 days. To start the mating step, THY.AP5 and THY.AP4 were transferred to YPD plates and grown for 1 d at 30°C. For the methionine assays, mated yeast was cultivated in selective liquid diploid medium for 3 d (30°C, 140 rpm). OD600 of selected yeast cells were measured and diluted to OD600 of 0.1 and 1.0 respectively in sterile water. Then 6 µL drops of each dilution were transferred to agar plates containing no (Met 0 µM) or 0.075 % (v/v) methionine (Met 50 µM). Subsequently, the yeast was grown for 72 h at 30°C and results documented photographically [103, 104].

### Subcellular localization and FRET measurements

The coding sequences of the genes of interest were fused in frame to yellow or cyan fluorescent protein as indicated and fluorescence resonance energy transfer (FRET)-experiments were performed as described by [105] with CLSM 780 (Zeiss). More than 20 protoplasts were measured twice for each set of PEG-based transformation and the experiment was repeated independently

## List of Abbreviation

ABA: abscisic acid
*A. thaliana*: *Arabidopsis thaliana*
AKIN10: protein kinase being part of the sucrose non-fermenting kinase complex
Asc: ascorbate
CaM: calmodulin
CML: calmodulin-like
DEG: differentially expressed gene
DHA: dehydroascorbate
DREB2A: Dehydration-responsive element binding protein 2
FC: fold change
GAPC: glyceraldehyde-3-phosphate dehydrogenase 2
GCN2: general control non-repressible 2
GO: gene ontology
GSH: glutathione
GSSG: diglutathione disulfide
HL: high-light (800 µmol photons m^−2^ s^−1^)
H_2_O_2_: hydrogen peroxide
HSE: heat shock element
HSF: heat shock factor
HSP: heat shock protein
IRES: independent ribosomal entry sites
JA: jasmonic acid
LL: low light (8 µmol photons m^−2^ s^−1^)
log_2_ FC: logarithmic fold change to base 2
MAPK: mitogen-activated protein kinase
mbSUS: mating-based Split-ubiquitin-System
MBF1c: Multiprotein Bridging Factor 1C
OPDA: 12-oxo-phytodienoic acid
ORF: open reading frame
PEG: polyethylene glycol
PET: photosynthetic electron transport
PTM: post-translational modification
RBP: RNA binding protein
ROS: reactive oxygen species
RuBisCO: ribulose bisphosphate carboxylase oxygenase
SAP: stress associated protein
SnRK1: SNF1Lrelated kinase 1
TE: translation efficiency
TF: transcription factor
TOR: target of rapamycin
TPT2: triose phosphate/phosphate translocator 2
uORF: upstream open reading frame
UTR: untranslated region
VTC2: GDP-L-galactose phosphorylase

## Acknowledgements

The authors acknowledge support by the Deutsche Forschungsgemeinschaft (Di 346/19; SP 1935) and for funding by the Australian Government through the Australian Research Council (DP220103640). Barry Pogson is the recipient of an Australian Research Council Australian Laureate Fellowship (FL190100056) funded by the Australian Government. Special thanks to Renate Scheibe for providing the pET16b construct for recombinant expression of GAPC1/GAPC2. We acknowledge the ACRF Biomolecular Resource Facility, a service node of Bioplatforms Australia, and the Centre for Biodiversity Analysis Ecogenomics and Bioinformatics Lab for provision of resources and expertise to perform Illumina sequencing. We acknowledge the facilities and the scientific and technical assistance of Microscopy Australia at the Centre for Advanced Microscopy, Australian National University, a facility that is funded by the University and the Federal Government.

The authors declare that they have no competing interests.

## Data availability

All relevant data is accessible in this manuscript or in the supplemental material. A preprint version is accessible on bioRxiv under: doi: https://doi.org/10.1101/2021.02.18.431817.

## Authors contributions

MM designed the research and performed the polysomal gradients, TEM microscopy, transcript profiling, microarrays, RPF-seq and total RNAseq and bioinformatic data analysis, transient protoplast reporter gene assays, RNA-EMSA, protein affinity pull downs and wrote the paper. AS did the RPF-library preparation and sequencing and performed bioinformatic analyses. CW prepared polysomal gradients and transcript profiling. SS performed Y2H experiments and cellular localization and FRET. MW performed RBP affinity pull downs and gel shift assays on SAP3 motifs. AF measured FRET and transcript profiles of OsSAP and OsCML49. DRG discussed and analyzed transcriptomics data. TS supervised and performed FRET measurements. BP supervised and discussed the results and wrote the paper. KJD designed the study, supervised and discussed the results, coordinated the group and wrote the paper.

## Supplementary materials

**Suppl. Fig. 1: Size distribution of RPF-Seq and frame analysis of start/end sites.** (a) The size distribution of RPF-sequencing reads from LL and LL➔HL-treated plants exhibited two distinct optima at 23-25 and 28-32 nucleotides matching the reported sizes for stalled plastidial/mitochondrial versus cytosolic footprints, respectively. (b,c) Periodicity of translation frames agrees with reported literature using combined application of cycloheximide and lincomycin in plant systems. Depicted are the mean normalized counts in RPKM for each coding frame (0, 1 or 2) at the relative nucleotide distance to the 3’- and 5’-end and start site, respectively. No difference in coverage could be detected between LL and LL➔HL conditions.

**Suppl. Fig. 2: Translationally regulated transcripts after 10 min LL➔HL.** (a) The volcano-plot of ribosome associated transcripts revealed 951 significantly (adj. p-value <0.05) changed transcripts after 10 min LL➔HL transfer. 76 % (793) of the significantly changed ribosome-associated transcripts were down-regulated with 15% (158) being upregulated and 8% (92) unchanged in their fold change response. (b) Transcriptional changes were less prevalent with only 204 total transcripts being significantly changed with 38% (83) down-regulated and 55% (121) upregulated total mRNA FC response. This corresponds to a 4.6-fold higher translational than transcriptional control of gene expression after 10 min LL➔HL transfer. (c,d) Despite being primarily categorized by ribosome association and not by altered transcription, cluster 2 is not distinguished when assessed using the by common translation efficiency (TE) calculations (TE = FC ribosome-associated mRNA / FC total mRNA). In fact, following TE calculation for this dataset (TE; as Ribo10/Total10 /Ribo 0/Total 0) confounds transcriptional and translational regulation and might incorporate different biological processes. For all TE-upregulated transcripts were the ribosome association (log_2_ Ribosomal mRNA FC (LL➔HL/LL)) and transcriptional changes (log_2_ Total mRNA FC (LL➔HL/LL)) plotted. The majority of TE upregulated transcripts are transcriptionally downregulated (95%; 456) with only 85% (404) of TE-upregulated transcripts actually having a positive fold change in ribosome association after 10 min LL➔HL transfer. More importantly, 15 % (73) of the TE-upregulated transcripts are less associated with ribosomes as is indicated by the downregulation of fold change in ribosome association, thus the group of TE upregulated transcripts consists out of positive association and negative association with ribosomes during stress.

**Suppl. Fig. 3: Cluster 1&2 representatives avoid global translational down-regulation in *akin10* and *mpk6* background.** (a-d) Four transcripts were selected as representatives from Cluster 1&2 and subjected to qPCR analysis in wildtype (WT), *mpk6* and *akin10* mutants. Data are means ± SE of n=5 independent experiments, * indicate significance difference relative to LL control, greek and capital letters indicate significant differences between genotypes for LL and LL➔HL samples, respectively. Two factor ANOVA with Tukey post hoc test: p <0.05, respectively. A significant increase of total transcript in LL was observed in *mpk6* and *akin10* in comparison to WT for *SAP2* as representative for Cluster 1 (a) and *SAP3* and *HSP70* as representatives for Cluster 2 (b,c), but not for *DREB2A* (d). Total transcript abundance of *SAP2* and *HSP70* was significantly higher in *mpk6* and *akin10* after LL➔HL-transfer than in WT, but not for *SAP3* and *DREB2A*. In *mpk6* under LL conditions, ribosome-associated transcript abundances were not significantly different for *DREB2A* and *SAP2* but significantly higher in *mpk6* than in WT for *SAP3* and *HSP70*. Ribosome-associated transcripts after LL➔HL transfer were significantly lower in *mpk6* than WT for *SAP2* and significantly higher for HSP70, with no significant difference for *DREB2A* and *SAP3*. In *akin10*, no significant difference to WT after LL➔HL for *SAP2* and *SAP3*, but a significant increased relative abundance for HSP70 and *DREB2A* ribosome-associated transcripts could be detected. The 10 min LL➔HL response was similar for all four ribosome-associated transcripts in mutants and WT with slightly changed amplitude of relative increase for *HSP70* and *SAP2*. (e,f) Differences between transcriptional and translational effects of mutants on LL and LL➔HL treated plants prompted us to conduct a principal component analysis using the qPCR data from (a-d) to query genotype-specific and treatment-specific effects. (e) The three different genotypes overlap in their median variances (large symbols) for the whole experiment, indicating that the variance in transcript response is independent on the genotype. (f) Including the treatment in the PCA analysis revealed no overlap between the median principal component between LL and LL➔HL transcripts, indicating that the treatment accounted for the variance in the transcriptional response

**Suppl. Fig. 4: Translated regions of representative transcripts from cluster 1 & 2.** Genome Browser (IGV) views of RPF-seq (green) and total RNA-seq (blue) abundance (read depth) for loci of representative transcripts from cluster 1 & 2. Tracks are colored according to sample/time-point (each track represents overlay of n=3 biological replicates). Black arrow indicate potential ribosomal stalling sites. Gene model is symbolized as blue box with introns represented as dashed line, CDS as bold, and 5’- and 3’UTR as thin line with arrows indicating read direction.

**Suppl. Fig. 5: Distribution and secondary structure of motif-carrying 5‘UTRs.** (a) Motif 1, 2 and 3 occurrences in their respective transcripts as revealed by MEME-suit. The secondary structure is given in Dot-Bracket notation and was calculated using the software *RNA Shapes Studio* [44], with Motifs drawn at corresponding site(s) for motif 1 (green), 2 (yellow) and 3 (red), respectively. (b) Distribution of motif-carrying transcripts in translatomes are indicated as dots for transcripts carrying motif 1 (green), 2 (yellow) or 3 (red).

**Suppl. Fig. 6: Reporter-construct analysis for translational regulatory elements and secondary structure.** (a) Motif 1, 2 and 3 occurrences in their respective transcripts as revealed by MEME-suit. The secondary structure is given in Dot-Bracket notation and was calculated using the software RNA Shapes Studio [44], with motifs drawn at corresponding site(s) for motif 1 (green), 2 (yellow) and 3 (red), respectively. (b) Distribution of motif-carrying transcripts in translatomes are indicated as dots for transcripts carrying motif 1 (green), 2 (yellow) or 3 (red).

**Suppl. Fig. 7: Motifs are potential RBP binding sites.** (a) 5‘-biotinylated M3-RNA with and without unlabelled competitor M3-RNA was incubated with total protein extracts of LL or 10 min LL➔HL shifted plants, respectively. Depicted is a representative blot. (b) RBP 1-D acrylamide gel. Biotinylated M1 and M2 RNAs were coupled to strep-tagged beads and incubated with protein extracts from LL treated plants. After washing, samples were eluted and loaded on one-dimensional acrylamide gels. The enriched bands (arrows) were cut out and analysed via MALDI. (c) RNA-EMSA was used to validate dose dependent interaction (ratio of bound to unbound) of *SAP3*-M1-RNA with arabidopsis cytosolic AtGapC1, AtGapC2 or GAPDH isolated from rabbit muscle GAPDH isoform C1 *in vitro*.

**Suppl. Fig. 8: Motifs 1-3 and SAP2, SAP3 translational regulation are conserved between *A. thaliana* and Rice.** OsSAP8 and OsSAP9 were identified as closest homologues of SAP2 and SAP3 from *A. thaliana*. (a) M1, M2 and M3 could be identified in the 5’-and 3’-UTR of the rice homologues with only slight variation. (b,c) qPCR-based quantification of *OsSAP8* and *OsSAP9* transcripts of total and polysomal RNA fractions after 10 min of L➔H-transfer confirms conserved translational regulation of SAP2 and SAP3 between *A. thaliana* and Rice. Data represent means ± SD of n=2 independent experiments, asterisks indicate significance relative to control, two sided Student‘s t-test: p <0.05.

**Suppl. Fig. 9: Subcellular localization of rice OsSAP8, OsSAP9, OsCML49a and OsSAP49b in transfected arabidopsis protoplasts and their molecular interaction shown by FRET.** (a) *A. thaliana* mesophyll protoplasts were transfected with OsSA8:YFP, OsSAP3:YFP, OsCML49b:CFP or OsCML49a:CFP vector DNA. Fluorescence images taken by CLSM for subcellular localization. (b) Following transfection with constructs, FRET efficiencies between CFP and YFP were quantified indicating conserved interaction between arabidopsis and rice SAP and CML49.

**Suppl. Fig. 10: Gene expression map of *AtSAP2*, *AtSAP3*, *AtCML49* and *OsSAP8, OsSAP9* and *OsCML49a*.** (a) Transcript abundance and expression pattern of *AtSAP2*, (b) *AtSAP3* and (f) *AtCML49* in tissues retrieved from the eFP Browser. (d) Transcript abundance and expression pattern of *SAP8*, (e) *SAP9* and (f) *CML49a* in different rice tissues retrieved from the Rice eFP Browser. *CML49b* was missing in group 1 data source of Rice eFP Browser.

**Suppl. Fig. 11: GO-Analysis of *CML49* & *SAP3* Co-Expressed Genes during HL Acclimation.** (a) List of co-regulated targets of SAP3 and CML49 that are stronger deregulated in *cml49* than in *sap3* background were subjected to GO-term analysis via Cytoscape BiNGO. Dark red nodes represent categories that are significantly overrepresented in context of genome abundance. The size of node represents gene abundance in the co-regulated gene set to the corresponding A strong overrepresentation of response to high light intensity, response to heat and response to hydrogen peroxide was detected.

---

**Suppl. Table 1.**
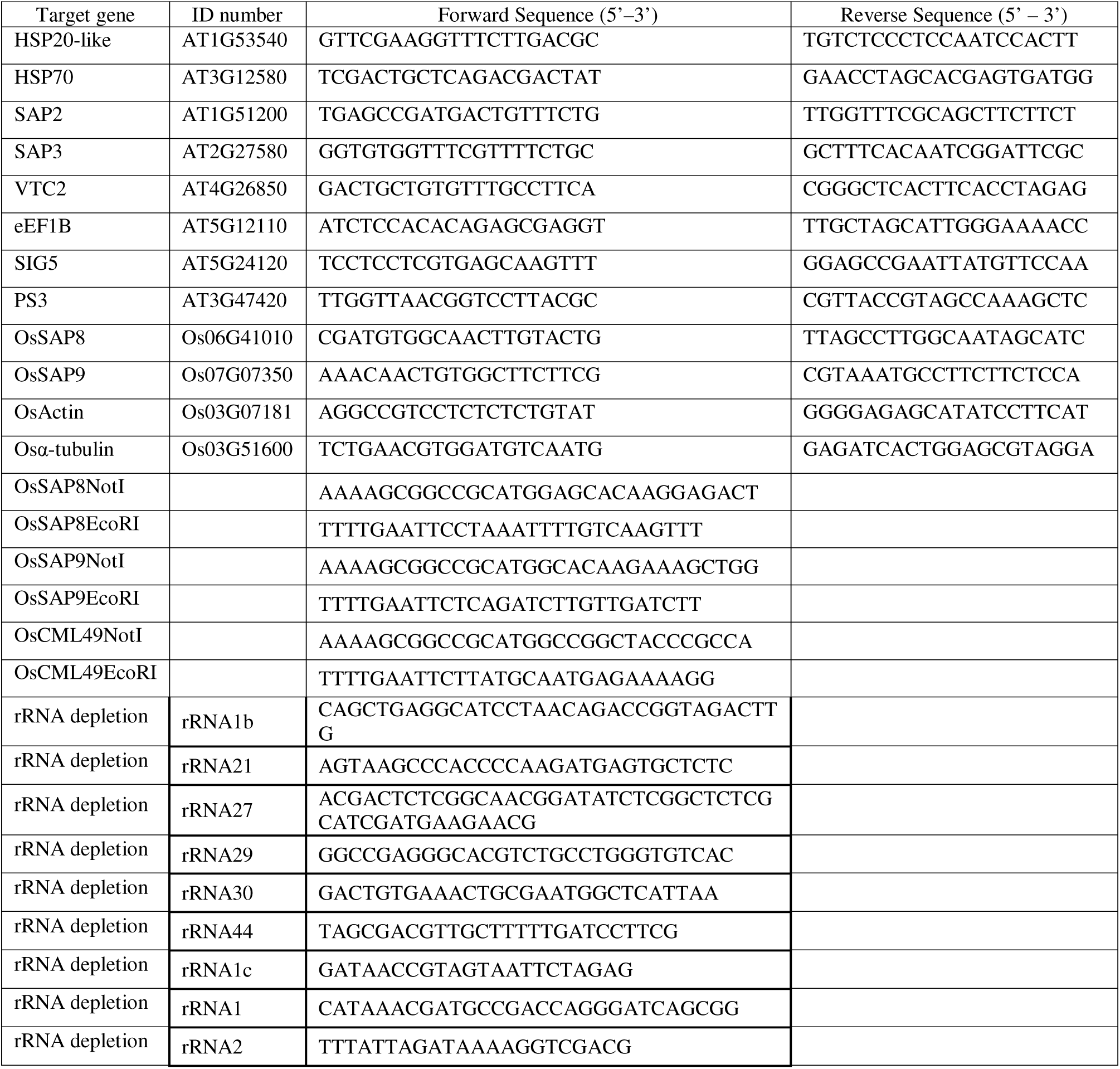
Oligonucleotides.

**Suppl. Table 2:** Overview of *cis*-elements within the promoter regions of *CML49* and *SAP3* co-regulated genes. The total number and type of *cis*-elements within the co-regulated subgroup was searched −1.000 bp upstream of the ATG region using PlantPAN2.0. The largest group of *cis*-elements belong to the WRKY-group, followed by MYB, MYB-related, SURE (sulfur-responsive element) and ABRE-like (early responsive to dehydration). Only few putative sites belong to the LS7 (salicylic acid-inducible element), C-box (bZIP factors), PHR1 (phosphate starvation response) and some represent cis-elements for light-dependent SORLIPs (Sequences Over represented in Light-Induced Promoters).

| AGI | Name | MYB/ MYB-related | WBOX | SURE | PHR1 | C-box | ABRElike | LS7 | SORLIPs | Other |
| --- | --- | --- | --- | --- | --- | --- | --- | --- | --- | --- |
| AT4G13800 | unknown | 29 | 37 | 35 | 4 | 2 | 20 | 4 | 17 | 24 |
| AT5G52570 | BETA-OHASE 2 | 24 | 36 | 29 | 6 | 2 | 27 | 4 | 30 | 26 |
| AT5G49480 | CP1 | 24 | 41 | 30 | 1 | 2 | 16 | 1 | 19 | 21 |
| AT4G04020 | FIB | 22 | 39 | 18 | 5 | 3 | 19 | 3 | 26 | 16 |
| AT5G58770 | CPT4 | 35 | 35 | 22 | 3 |  | 23 | 3 | 30 | 29 |
| AT3G56290 | unknown | 23 | 47 | 31 | 3 | 2 | 25 | 3 | 35 | 34 |
| AT1G67360 | REF | 41 | 46 | 32 | 5 | 8 | 30 | 6 | 27 | 14 |
| AT1G61800 | GPT2 | 28 | 36 | 23 | 7 | 9 | 18 | 3 | 17 | 26 |
| AT5G20150 | SPX1 | 22 | 42 | 20 | 13 | 4 | 14 | 1 | 10 | 23 |
| AT3G21890 | BBX31 | 22 | 41 | 27 | 2 | 6 | 22 | 3 | 14 | 30 |
| AT2G46830 | CCA1 | 31 | 43 | 47 | 1 | 6 | 21 | 1 | 44 | 14 |
| AT5G52640 | HSP90.1 | 35 | 38 | 43 | 4 | 4 | 23 | 3 | 15 | 14 |
| AT2G36800 | DOGT1 | 32 | 48 | 44 | 5 | 6 | 28 | 3 | 30 | 14 |
| AT5G48570 | ROF2 | 24 | 49 | 30 | 2 | 4 | 22 | 1 | 15 | 14 |
| AT1G52690 | LEA7 | 27 | 44 | 27 | 5 | 8 | 32 | 4 | 28 | 25 |
| AT2G24540 | AFR | 28 | 45 | 41 | 6 | 9 | 28 | 4 | 28 | 31 |
| AT3G09350 | Fes1A | 34 | 41 | 40 | 3 | 9 | 26 | 9 | 25 | 21 |
| AT4G21990 | APR3 | 22 | 40 | 26 | 3 | 6 | 21 | 1 | 16 | 20 |
| AT4G12400 | HOP3 | 33 | 37 | 25 | 3 | 1 | 24 | 6 | 12 | 21 |
| AT5G50800 | ATSWEET13 | 26 | 43 | 32 | 4 | 7 | 24 | 5 | 30 | 19 |
| AT2G37180 | RD28 | 23 | 43 | 21 | 7 | 4 | 22 | 3 | 20 | 34 |
| AT4G34550 | PCH1-like | 26 | 35 | 30 | 4 | 3 | 16 | 3 | 19 | 34 |
| AT3G17609 | HYH | 27 | 38 | 27 | 6 | 5 | 24 | 3 | 28 | 30 |
| AT5G64510 | TIN1 | 34 | 44 | 34 | 4 | 12 | 27 | 9 | 30 | 30 |
| AT1G01520 | ASG4 | 33 | 34 | 45 | 3 | 2 | 29 | 7 | 38 | 26 |
| AT1G32900 | GBSS1 | 31 | 52 | 31 | 5 | 9 | 28 | 2 | 36 | 30 |
| AT2G26150 | HSFA2 | 20 | 60 | 46 | 0 | 4 | 19 | 3 | 30 | 34 |
| AT5G12110 | eEF1B | 41 | 53 | 34 | 2 | 6 | 17 | 4 | 32 | 31 |
| AT4G26850 | VTC2 | 20 | 40 | 31 | 8 | 8 | 23 | 4 | 20 | 36 |
| AT2G33380 | RD20 | 29 | 42 | 26 | 6 | 7 | 38 | 3 | 22 | 29 |
| AT3G21150 | BBX32 | 26 | 41 | 39 | 5 | 6 | 17 | 5 | 30 | 23 |
| AT3G09450 | unknown | 38 | 34 | 42 | 6 | 1 | 29 | 3 | 43 | 29 |
| AT2G21320 | BBX18 | 26 | 57 | 40 | 7 | 4 | 20 | 1 | 20 | 27 |
| AT1G58520 | RXW8 | 20 | 28 | 26 | 2 | 3 | 17 | 4 | 18 | 45 |
| AT5G24120 | SIG5 | 14 | 52 | 31 | 1 | 4 | 22 | 1 | 16 | 25 |
| AT5G65380 | MATE | 24 | 47 | 31 | 4 | 8 | 32 | 7 | 33 | 29 |
| AT3G53830 | RCC1 | 24 | 40 | 35 | 6 | 4 | 31 | 2 | 18 | 27 |
| AT3G17790 | PAP17 | 37 | 44 | 34 | 7 | 8 | 28 | 6 | 31 | 29 |
| AT1G54050 | HSP20-like | 44 | 40 | 30 | 6 | 5 | 15 | 3 | 38 | 31 |
| AT1G62180 | ATAPR2 | 29 | 53 | 24 | 5 | 3 | 26 | 3 | 27 | 28 |
| AT3G21670 | ATNPF6.4 | 22 | 44 | 27 | 3 | 1 | 25 | 2 | 21 | 37 |
| AT3G12580 | HSP70 | 20 | 42 | 24 | 4 | 1 | 18 | 2 | 11 | 32 |
| AT2G46790 | PRR9 | 22 | 43 | 26 | 6 | 3 | 19 | 1 | 44 | 33 |
| AT1G77450 | NAC032 | 31 | 45 | 35 | 4 | 6 | 33 | 8 | 35 | 30 |
| AT2G15020 | unknown | 20 | 34 | 26 | 5 | 0 | 24 | 3 | 34 | 36 |
| AT3G47420 | PS3 | 30 | 42 | 10 | 7 | 2 | 21 | 5 | 18 | 27 |
| AT5G63130 | Phox/Bem1p | 20 | 49 | 19 | 4 | 5 | 15 | 3 | 31 | 37 |
| AT1G01720 | ANAC2 | 29 | 39 | 50 | 2 | 6 | 22 | 2 | 36 | 23 |
| AT4G26150 | CGA1 | 26 | 34 | 27 | 5 | 2 | 15 | 2 | 14 | 36 |
| AT1G74310 | HSP101 | 20 | 35 | 33 | 5 | 2 | 13 | 1 | 17 | 34 |
| Total |  | 1368 | 2112 | 1556 | 224 | 232 | 1148 | 173 | 1278 | 1368 |

**Suppl. Table 3.**
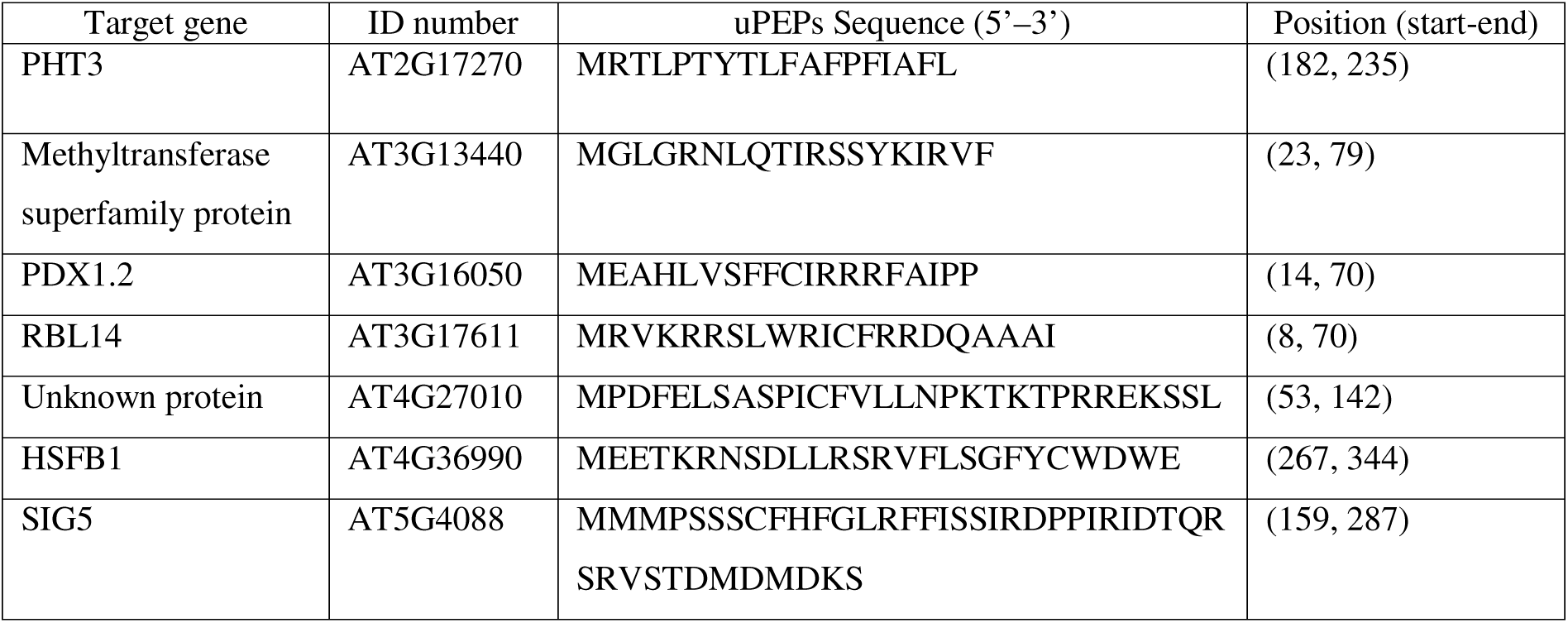
uORF analysis of motif-carrying transcripts. The motif-carrying transcripts were screened for upstream open reading frames (uORF) coding for upstream peptides (uPEPs) using the online tool uPEPperoni.

