## Supplemental Figures for "Retrograde Control of Cytosolic Translation Targets Synthesis of Plastid Proteins and Nuclear Responses for High-Light Acclimation"

Suppl. Figure 1

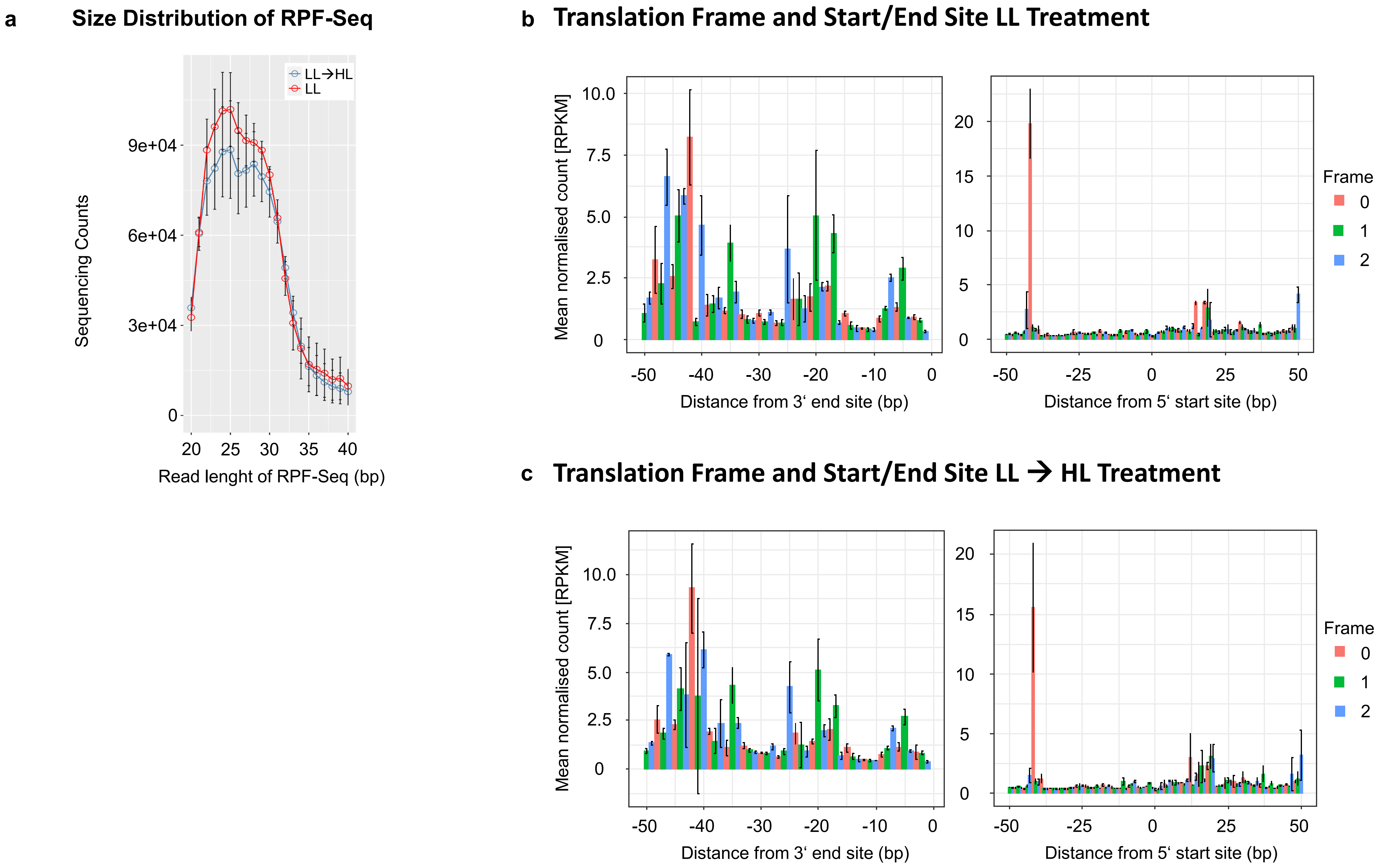

**Suppl. Figure 2**

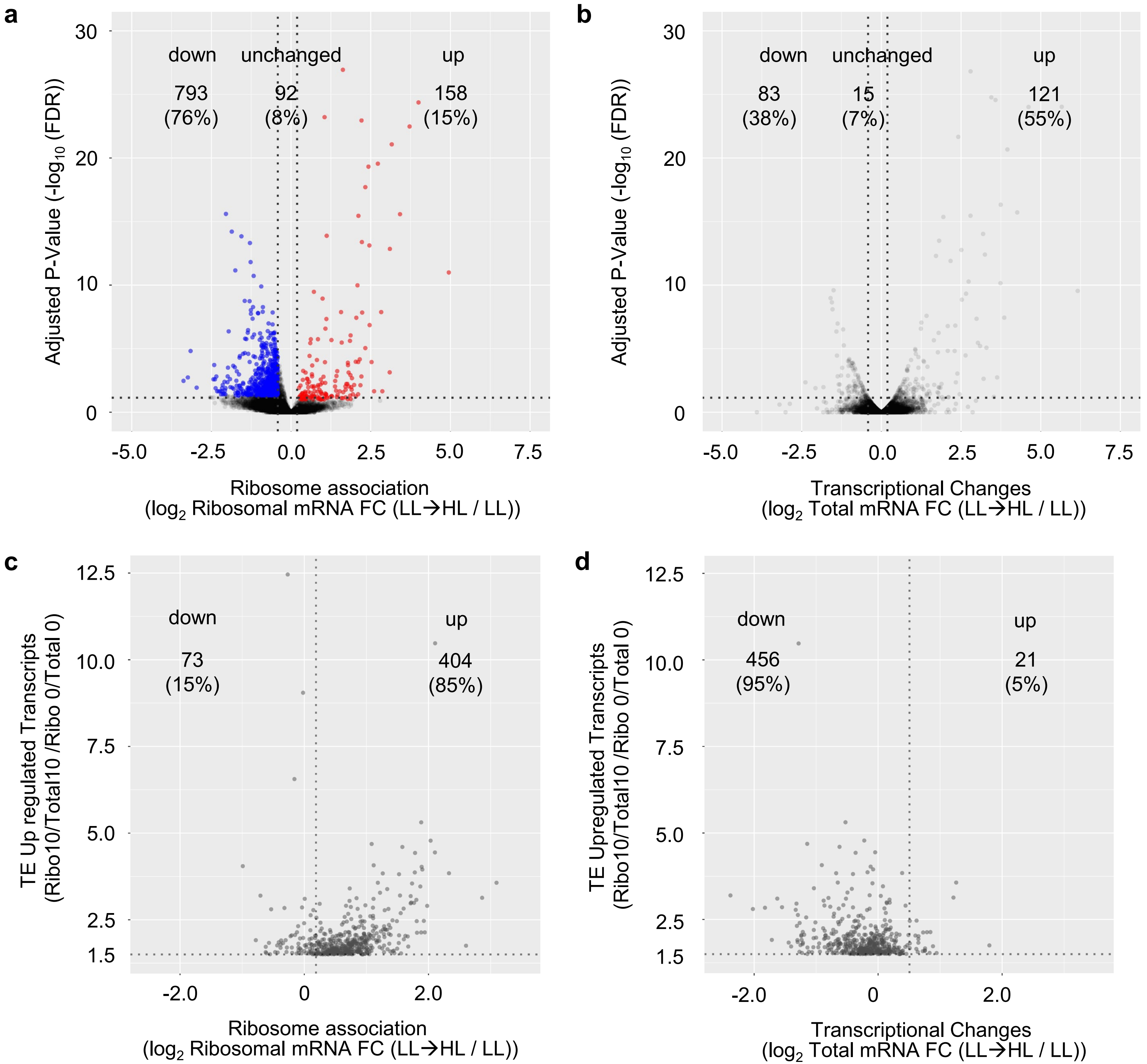

**Suppl. Figure 3**

#### Cluster 1&2 representatives avoid cumulative translational down regulation in *akin10* and *mpk6* background

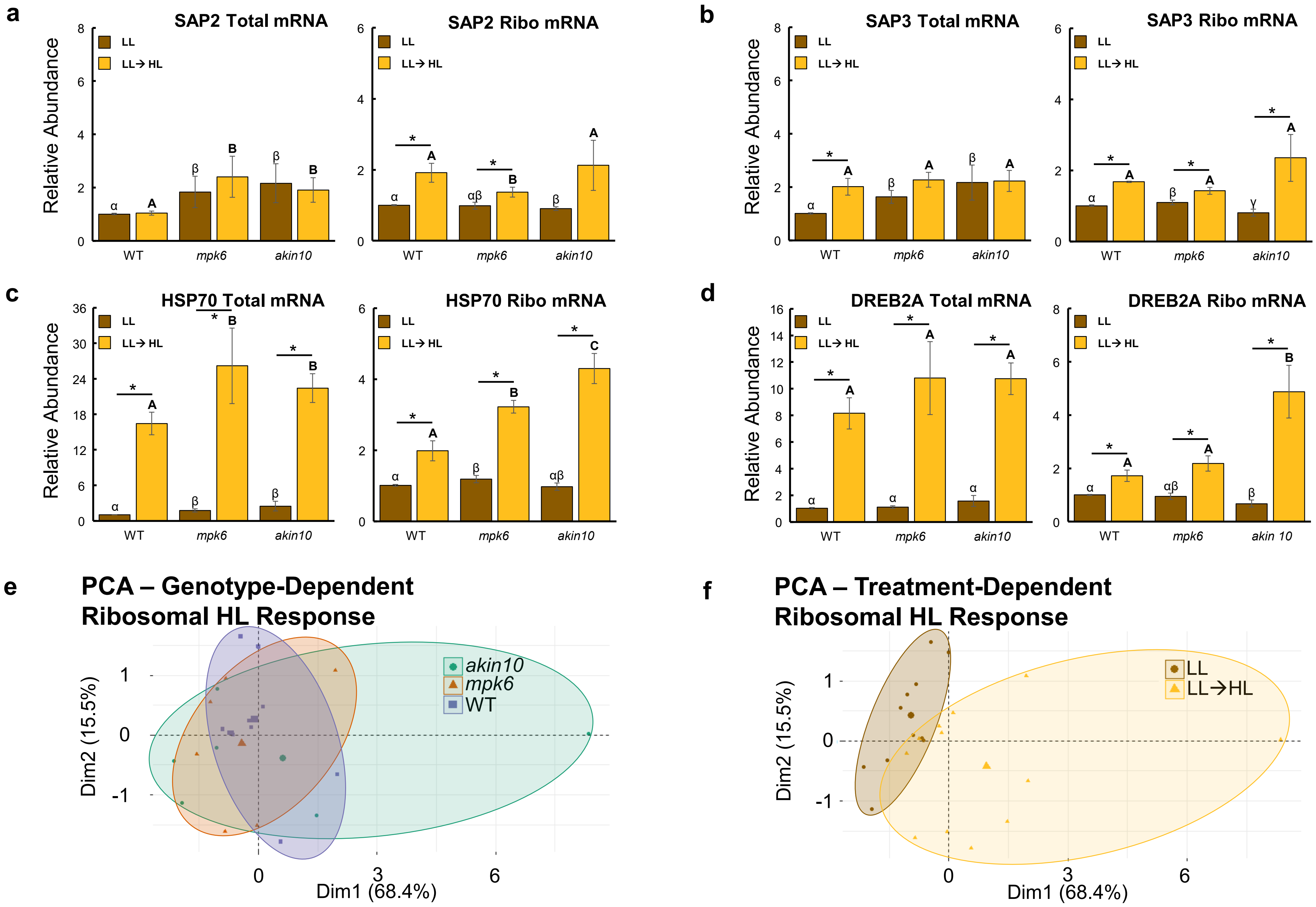

**Suppl. Figure 4**

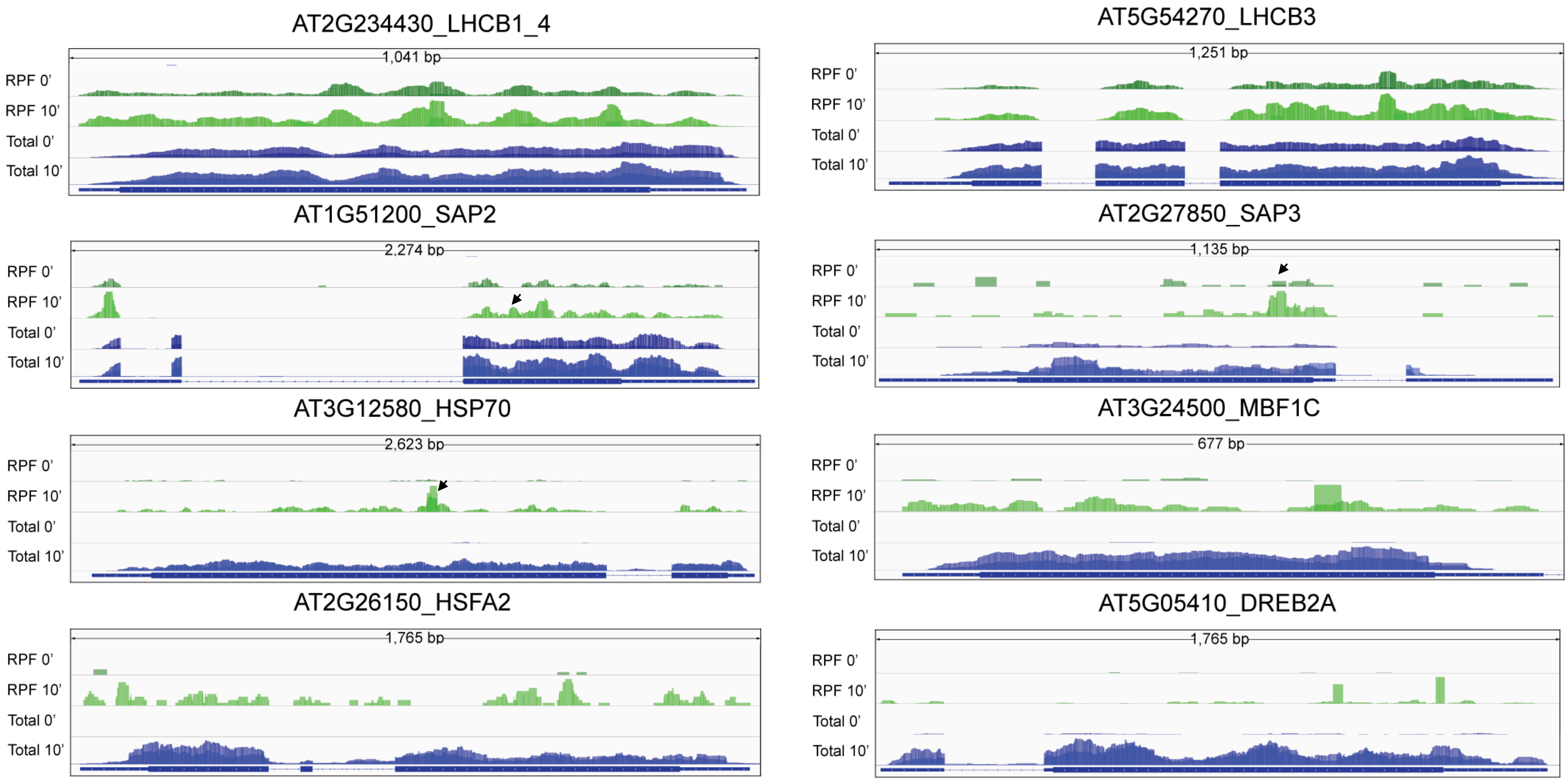

**Suppl. Figure 5**

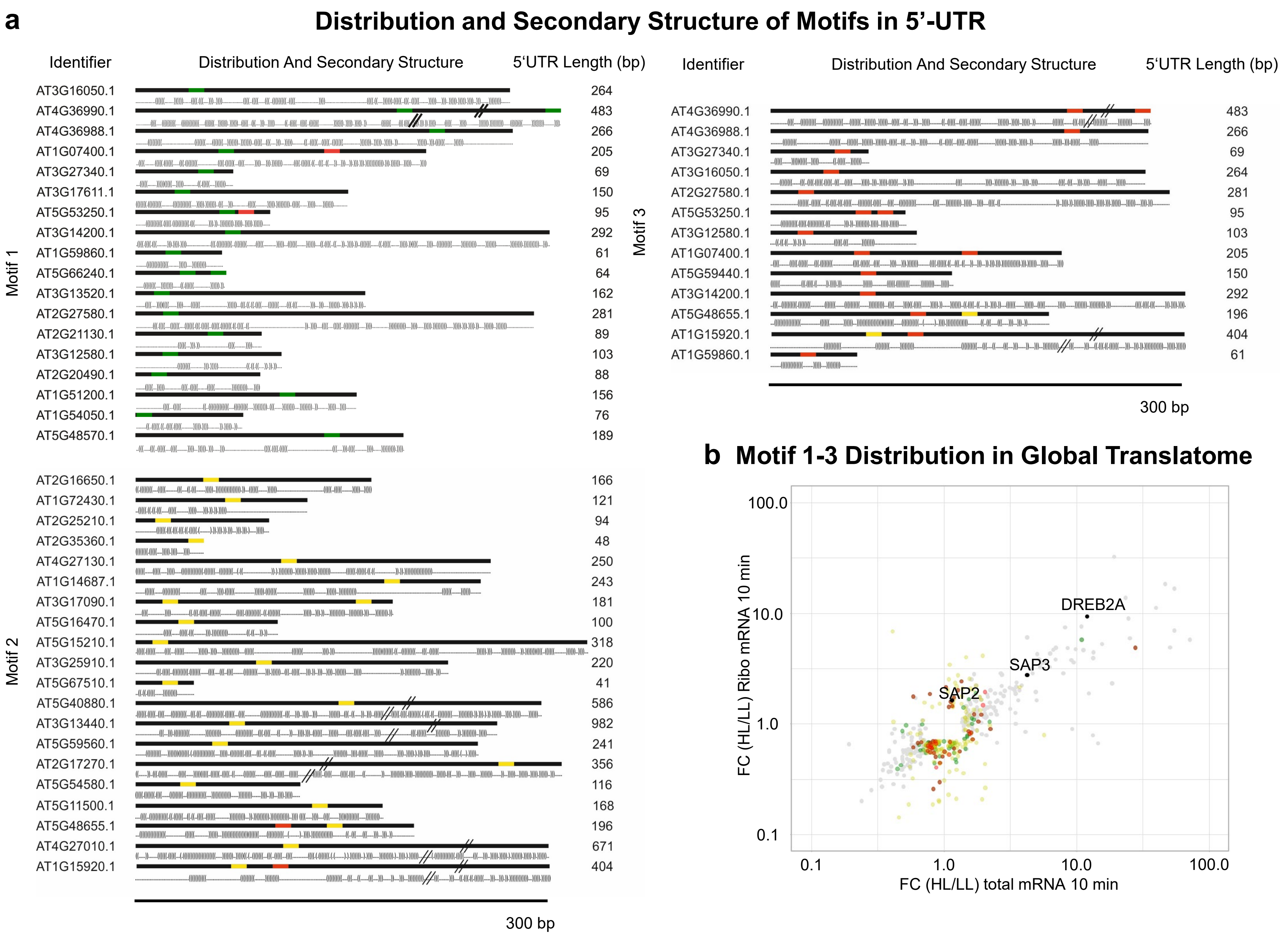

**Suppl. Figure 6**

#### Reporter-Construct Analysis for Translational Regulatory Elements and Secondary Structure

| 5'-UTR Construct | δE | p value | Secondary Structure (Dot-Bracket) | | Secondary Structure Model |
|---|---|---|---|---|---|
| **Empty Ubi** | -13.64 | 0.039 | ........(((((.......))))).................................(((((.......))))).....  | [ ][ ] | 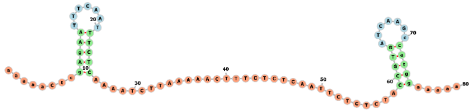 |
| **Motif 1** | -41.52 | 0.034 | ........(((((((................((((((((((...)))))))))).....(((((((.....))))))).)))))))....(((((.......))))).....  | [ [ ] [ ] ] [ ] | 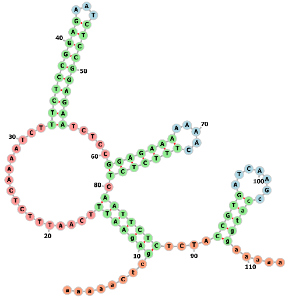 |
| **Motif 2** | -41.68 | 0.023 | ..((((((((((.......)))))..((((....)))).))))))((((...(((((.((((...(((((....)))))..))))))))))))..(((((.......))))).....  | [ [ ] [ ] ] [ ] [ ] | 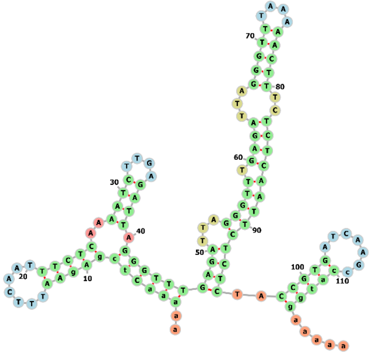 |
| **Motif 3** | -32.53 | 0.011 | ........(((((........)))))......((((..(((.....)))(((.....(((((((...))))))).......)))......((((.......))))))))..  | [ ] [ [ ] [ ] ] [ ] ] | 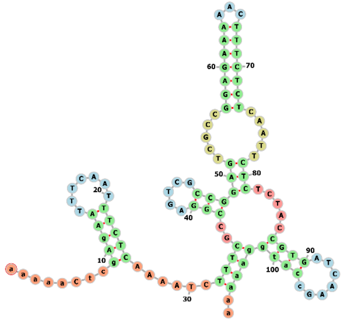 |

* Red marker for:
EGFR Sequence (epidermal growth factor receptor) – 1st Intron downstream enhancer;
T Maekawa, F Imamoto, G T Merlino, I Pastan, S Ishii (1989)

### Suppl. Figure 7

#### Motif3 RNA Protein Pulldown and Identification

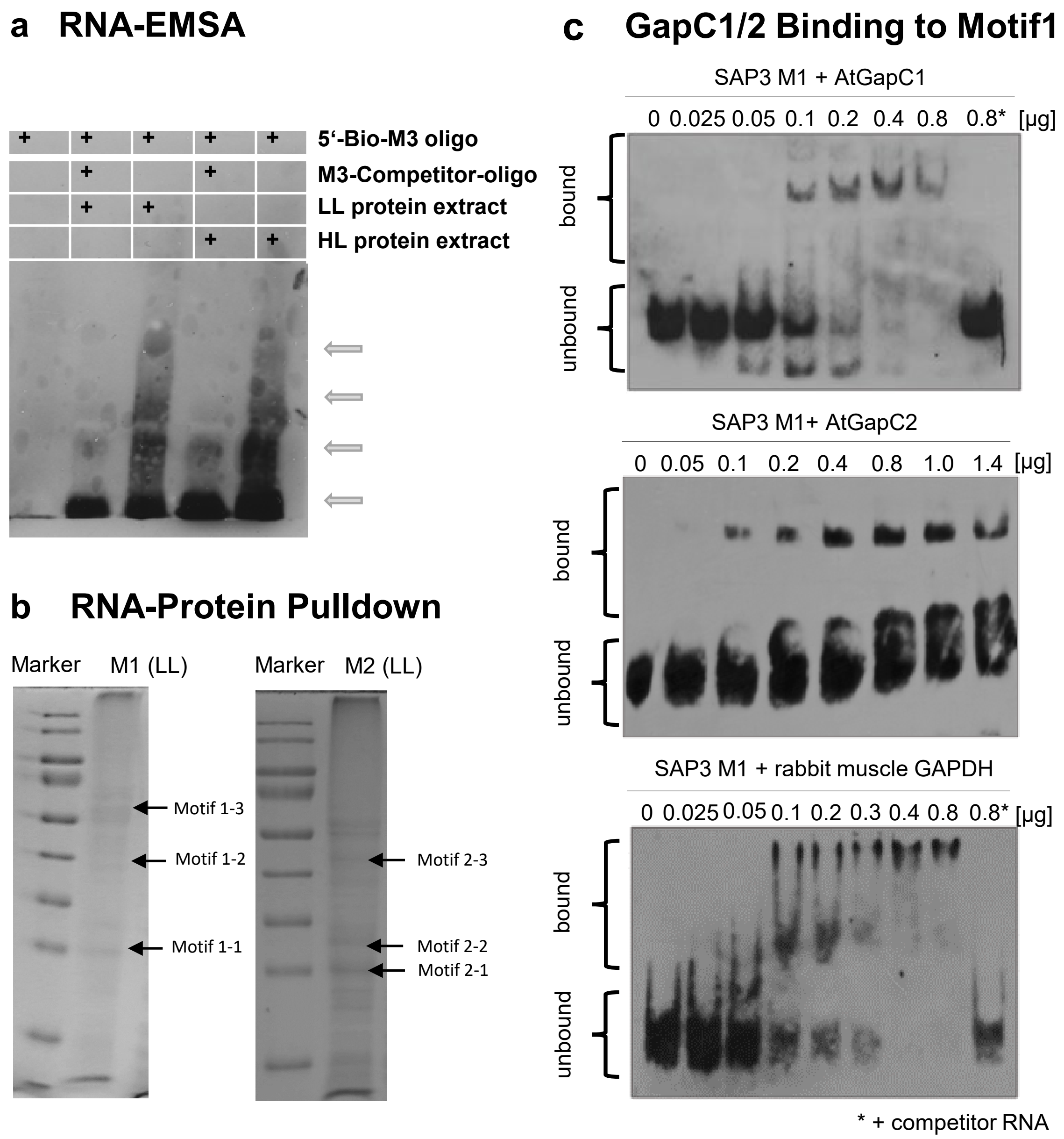

**Suppl. Figure 8**

**a Enriched Motifs are Conserved across Species**

***Arabidopsis motifs***

***Rice motifs***

**Motif 1**

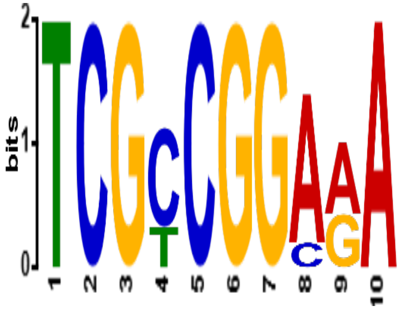


**OsSAP8:**
**TCGCCGGAGAG**
**OsSAP9:**
**TCTCGGGAGAA**

**Motif 2**

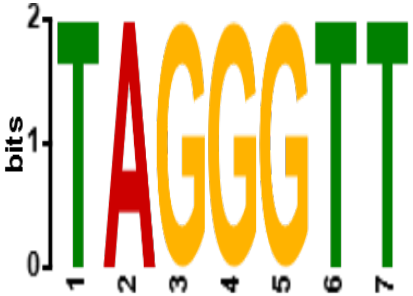


**OsSAP8:**
**TAGGGTTC**
**OsSAP9:**
**AGAGGTTT**

**Motif 3**

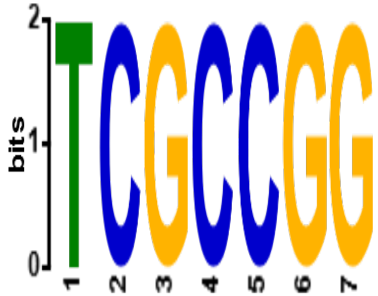


**OsSAP8:**
**TCGCCGGAG**
**OsSAP9:**
**CCTCCGGTG**

**b Translational Control of Rice SAP2/3 Homologues**

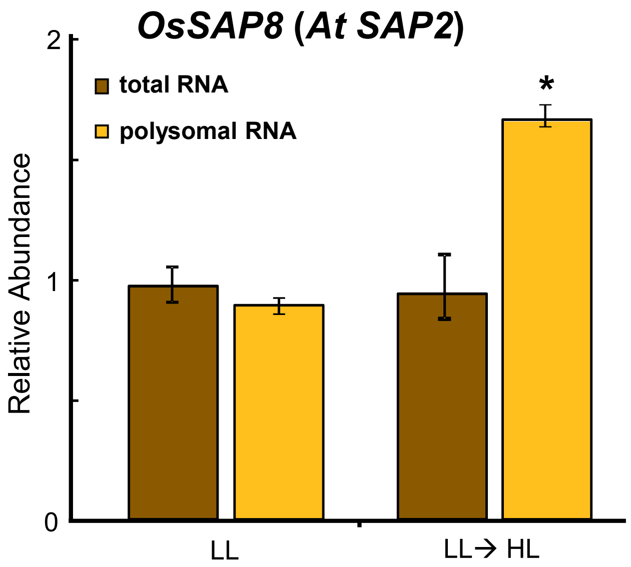


**c**

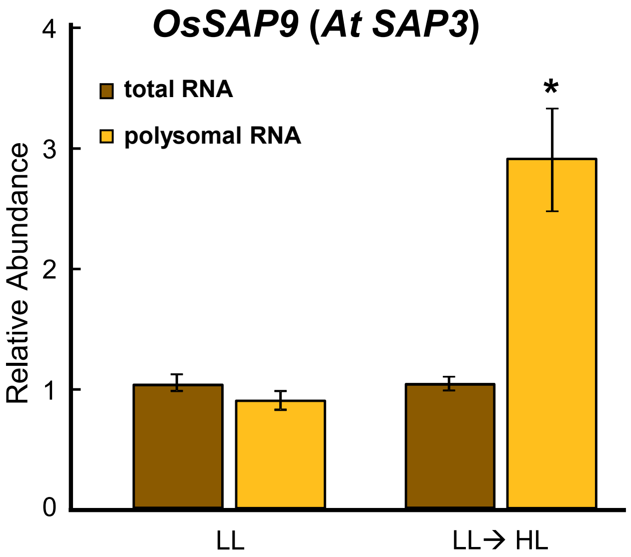

#### Suppl. Figure 9

**Conserved SAP2 and SAP3 Interaction and Re-Location with CML49**

**a** ***In-vivo*** **Co-Localization of OsSAP8, OsSAP9 and OsCML49a/b**

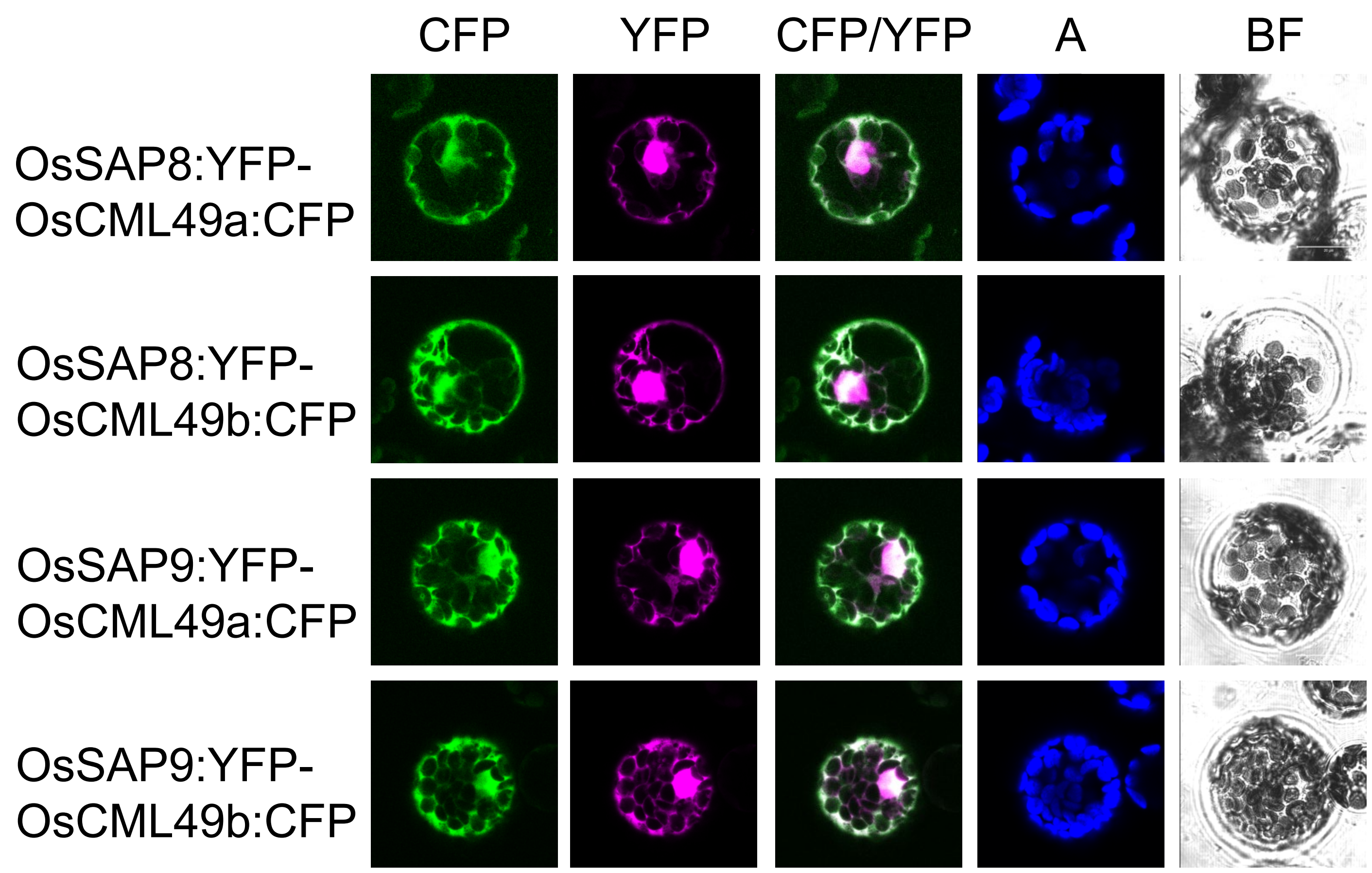


**b** **FRET Interaction of OsSAP8, OsSAP9 and OsCML49a/b**

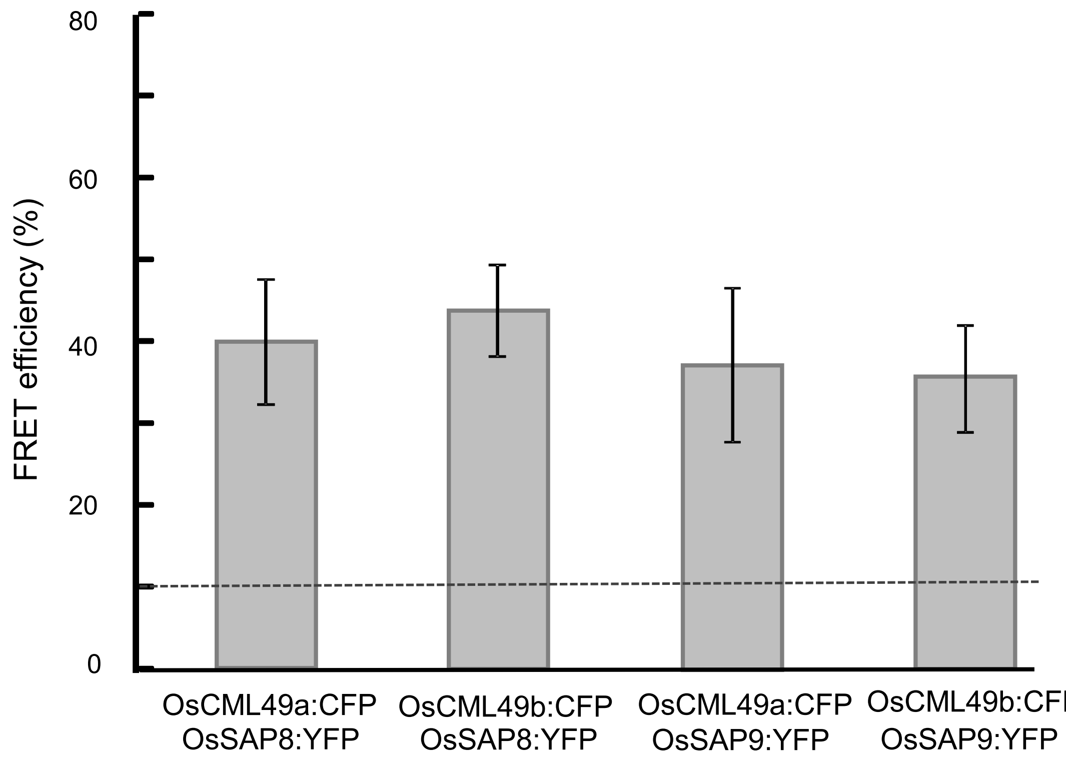

### Suppl. Figure 10

#### Expression Pattern of SAP2/3 and CML49 Homologues in *Arabidopsis* and Rice

##### a SAP2 Expression Pattern

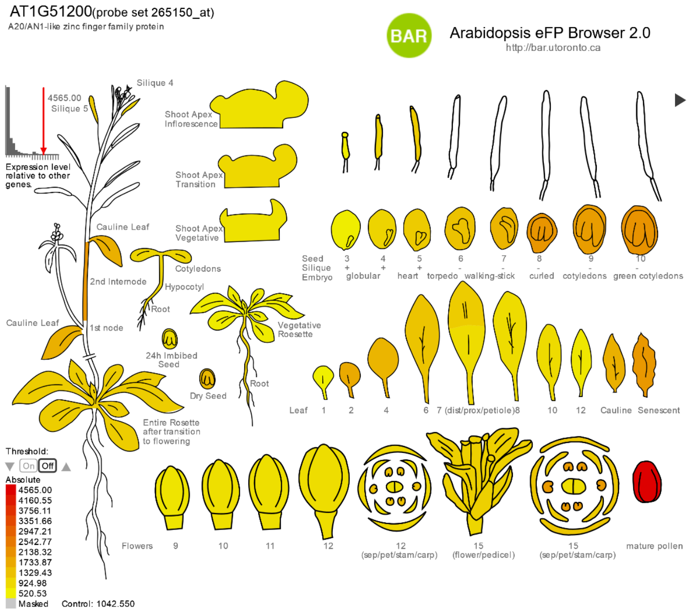


##### b SAP3 Expression Pattern

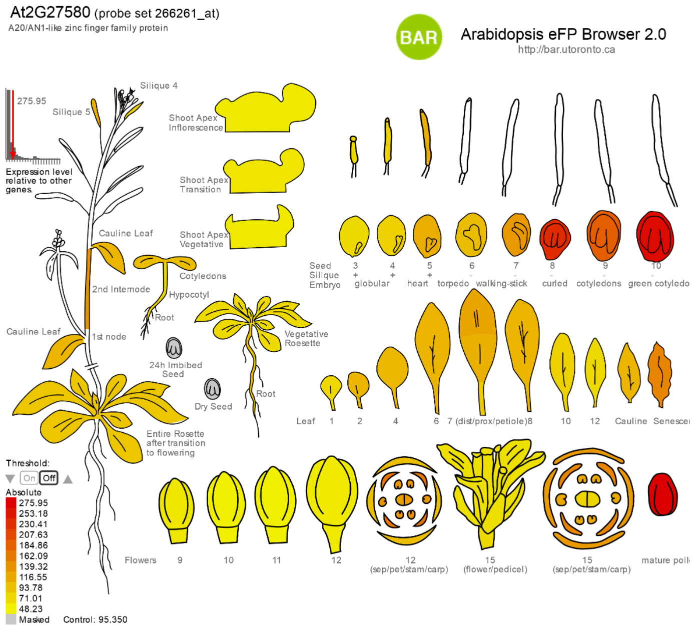


##### c CML49 Expression Pattern

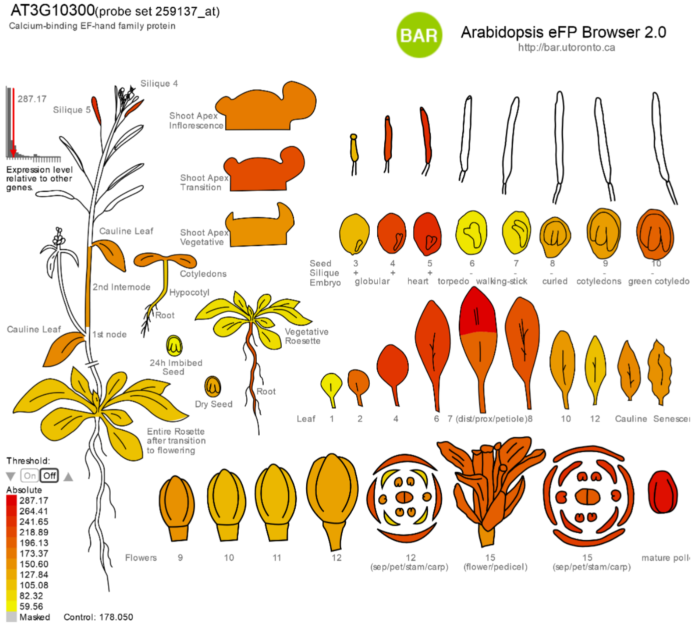


##### d OsSAP8 Expression Pattern

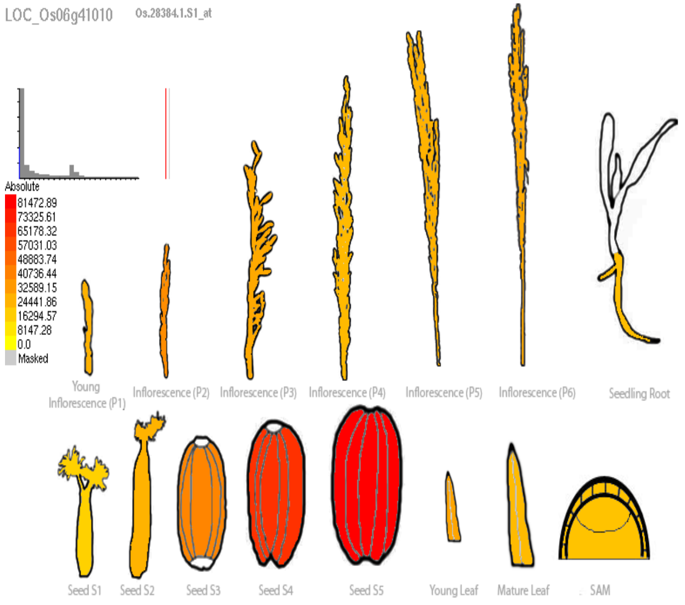


##### e OsSAP9 Expression Pattern

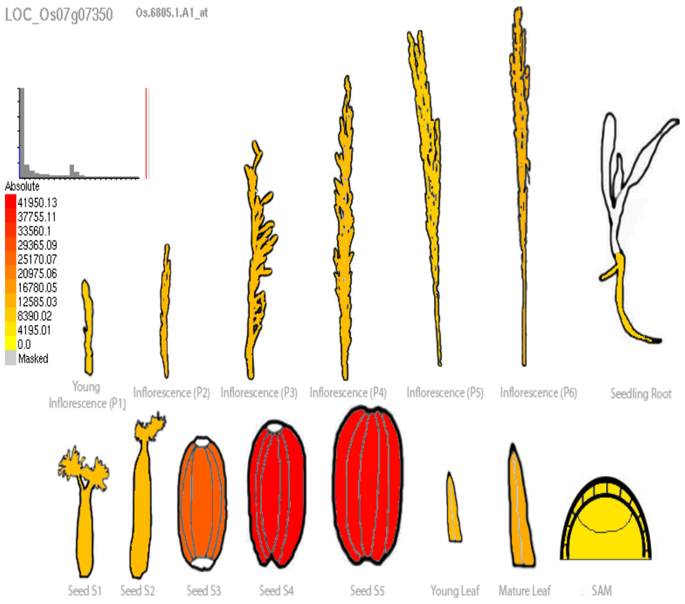


##### f OsCML49 Expression Pattern

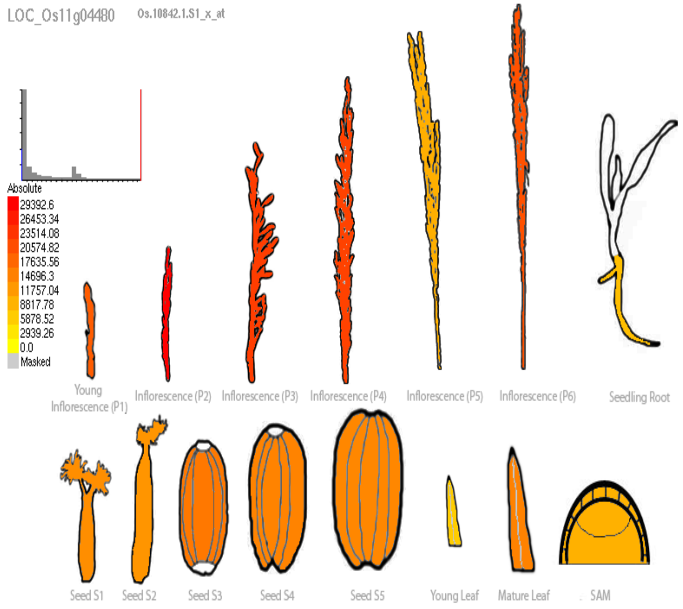

**Suppl. Figure 11**

**GO-Analysis of CML49 & SAP3 Co-Expressed Genes during High Light Acclimation**

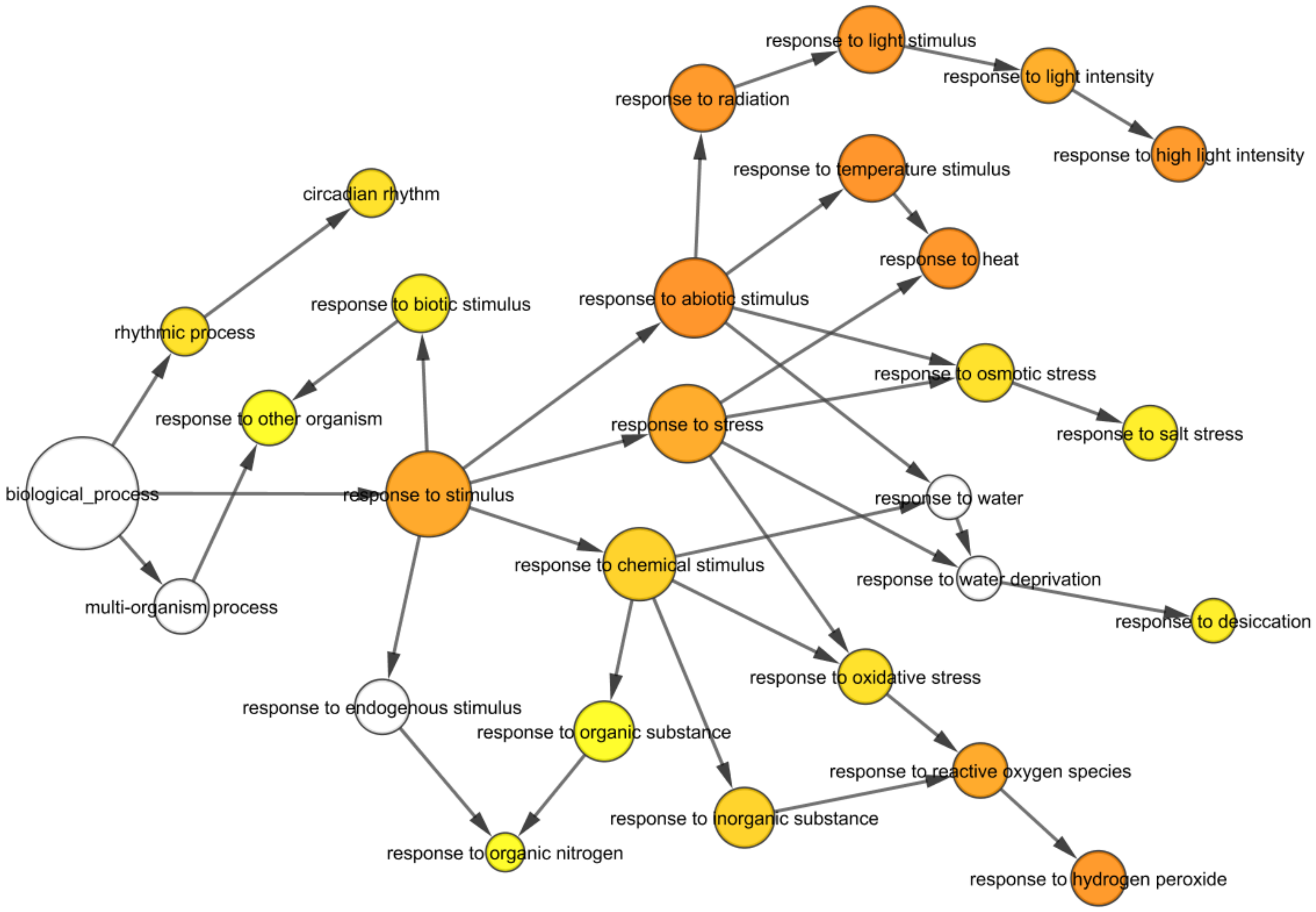
